# R-CHOP in B cell non-Hodgkin lymphoma: balancing anti-tumor efficacy and NK-cell functionality

**DOI:** 10.64898/2026.01.12.698321

**Authors:** Johanna Jansky, Hannah Breitung, Sylvia Zöphel, Laetitia Döngi, Florian Wentzel, Daniel Picard, Nadja Küchler, Hermann Eichler, Marc Remke, Moritz Bewarder, Lorenz Thurner, Gebhard Stopper, Eva C. Schwarz, Markus Hoth

## Abstract

Combining the anti-CD20 antibody rituximab with the chemotherapeutic drugs vincristine, doxorubicin, and cyclophosphamide plus the corticosteroid prednisone (R-CHOP) has been the standard first line-line therapy for many non-Hodgkin lymphomas including diffuse large B-cell lymphoma (DLBCL) for more than 20 years. Natural killer (NK) cells are considered major mediators of rituximab-induced antibody-dependent cellular cytotoxicity (ADCC). While the anti-tumor effects of vincristine, doxorubicin, and cyclophosphamide are well-documented, their impact on NK cell viability and effector function remains poorly characterized.

We evaluated the single-agent and combinatorial efficacy of vincristine, doxorubicin, and the cyclophosphamide surrogate mafosfamide in four DLBCL cell lines (TMD8, WSU-DLCL2, U-2932, and RI-1) relative to their impact on NK cell survival and effector function. In single-agent experiments, vincristine eradicated all B-cell lines in a dose-dependent manner at clinically relevant concentrations, whereas doxorubicin and mafosfamide required higher doses to achieve comparable efficacy. Although all three agents reduced NK cell viability over time at clinically relevant concentrations, impaired NK cell cytotoxicity was observed only at supra-clinical doses. Combinatorial experiments revealed antagonistic interactions between vincristine and doxorubicin, without evidence of significant synergistic effects. Incorporating tumor and NK cell survival into a mathematical model support a measurable trade-off between tumor cell reduction and immune cell viability. To assess the translational relevance of our *in vitro* findings, we analyzed a cohort of 32 patients with B-cell non-Hodgkin lymphoma. We confirmed NK cell depletion during R-CHOP and R-mini-CHOP therapy, however NK cell loss was significantly less pronounced in patients receiving dose-reduced R-mini-CHOP. Notably, the intrinsic cytotoxicity of residual NK cells was preserved under both treatment regimens, with a trend toward enhanced cytotoxicity during R-mini-CHOP treatment. These findings suggest that R-CHOP predominantly affects NK cell abundance rather than function and support the translational relevance of the *in vitro* results.

## Introduction

Diffuse large B-cell lymphoma (DLBCL) is the most commonly diagnosed type of non-Hodgkin lymphoma (NHL), accounting for 25-35% of all cases (2), with a prevalence of about 7 cases per 100,000 people per year (3). DLBCL is thus the (or one of the) most common blood cancers. It is a heterogeneous disease, with around 85% of cases falling into one of two subgroups: germinal centre B-cell-like (GCB) and activated B-cell-like (ABC). These subgroups are defined by similarities in gene expression to the cell of origin (2).

The era of modern chemotherapy probably began in 1931 with the treatment of cancer by dichlorethylsulphide (mustard gas) (4) as discussed in more detail in the review by Dy and Adjei (5). As early as 60 years ago, lymphomas and other blood cancers were already being treated with chemotherapeutic agents like cyclophosphamide (6), vincristine (7) or doxorubicin (8). These and other drugs were often combined to treat different types of lymphoma. For example, cyclophosphamide and vincristine were combined with the corticosteroid prednisone in a regime known as COP already 50 years ago (9, 10). Combinatorial drug therapies are believed to be superior to monotherapies because they are less likely to result in resistance. In the seventies, bleomycin, doxorubicin, cyclophosphamide, vincristine and prednisone (BACOP) were combined to treat advanced non-Hodgkin lymphomas (11, 12). At the same time, cyclophosphamide, doxorubicin (hydroxydaunorubicin), vincristine and prednisone (CHOP) were combined to treat malignant lymphoma (13). When compared with other treatment regimens, CHOP finally became the preferred treatment for many forms of NHL including DLBCL (14–17).

The human B lymphocyte specific antigen CD20 (formerly also known as B1) was identified by monoclonal antibodies (18). CD20 is the target for rituximab, which was the first antibody approved by the FDA (1997) and the EU (1998) for the treatment of cancer (19–21). It is probably not surprising that CHOP and rituximab were eventually combined to create R-CHOP for treatment of DLBCL and other NHLs (22–26).

R-CHOP therapy has been the standard first-line therapy until today. It is usually administered in six to eight cycles, with the number of cycles or dose intensity sometimes being modified for certain age or risk groups. In 2022, the results of a clinical trial combining polatuzumab vedotin, rituximab, cyclophosphamide, doxorubicin and prednisone (pola R-CHP) were published (27). This combination was approved by the FDA as a first-line therapy for adult patients in 2023, providing an alternative to R-CHOP. Very recent results describe a slight benefit of pola R-CHP vs R-CHOP in patients with previously untreated intermediate- or high-risk DLBCL (28).

In addition to R-CHOP, pola R-CHP and their modifications, significant breakthroughs have been achieved for patients with relapsed or refractory DLBCL, even without chemotherapy or with chemo-light regimens (see (29) for an overview). Several approaches are now available as options for patients with relapsed or refractory DLBCL, including CAR (chimeric antigen receptor) T cells, as indicated by several promising studies (30–33). Additionally, glofitamab, a bispecific antibody recently approved by the FDA and the EU, which links T cells to (lymphoma) B-cells (34, 35), is a promising therapeutic tool in combination with gemcitabine and oxaliplatin (GemOx) for patients with relapsed or refractory DLBCL (35–37).

Natural killer cells (NK cells) play a vital role in the therapeutic approach against non-Hodgkin lymphomas including DLBCL, as they are thought to be crucial mediators of rituximab-induced cytotoxicity in B-lymphoma cells during R-CHOP or pola R-CHP therapy. Rituximab binds to CD16, which is mainly expressed on NK cells. It thereby couples endogenous NK cells (via CD16) and B-lymphoma cells (via CD20) and induces antibody-dependent cellular cytotoxicity (ADCC) of NK cells against B-lymphoma cells. It is believed that NK cells are the prime target of rituximab therapy, and it has been shown that NK cell numbers correlate with clinical outcome for NHLs (38, 39). This view has recently been challenged by findings indicating that the number of CD16+ T cells and CD16+ monocytes in peripheral blood is a more precise indicator of the outcome of R-CHOP treatment in patients with DLBCL than the number of CD16+ NK cells (40).

Considering that R-CHOP has been the standard therapy for several types of B-cell NHL including DLBCL, for many years, it is somewhat surprising that only few papers have been published on the optimization of the dosage of the three R-CHOP drugs cyclophosphamide, doxorubicin and vincristine (41). Furthermore, few studies have examined plasma concentrations in patients (42). According to our literature search, only limited data has been published on the potential side effects on NK cells or other immune cells. However, immune cell viability and functionality are fundamental to rituximab-based therapy. In our opinion, future clinical studies would benefit from simultaneously quantifying the effects of cyclophosphamide, doxorubicin and vincristine on different B-lymphoma cells (cell lines and/or primary tumor cells), as well as their impact on NK cell viability and cytotoxicity against the respective cancer cell lines. This could help to personalize future multimodal treatments for non-Hodgkin lymphomas including DLBCL. It could also serve as blueprint for studying the impact of chemotherapy on various immune cell compartments, such as T cells, given that current and future clinical trials increasingly incorporate different immunotherapy combinations. T cell engaging therapies like bispecific antibodies are currently studied in phase 3 trials in combination with CHOP-like chemotherapy (43–45). However, the exact effects of each chemotherapy component on T cell function remain unknown. As new immunotherapeutic options emerge, de-escalation strategies that omit chemotherapy cycles or individual components are being tested more frequently in clinical trials (46). Our results could also help inform decisions about which chemotherapy components may be omitted when combined with different kinds of immunotherapy, when designing and planning new clinical trials.

## Results

To investigate the effects of cytotherapeutic drugs *in vitro*, we incubated DLBCL cell lines and expanded NK cells with various concentrations of the respective chemotherapeutic agents, individually and in combination, for 24, 48 and 72 hours. The incubation doses were based on the plasma levels of DLBCL patients shortly after drug administration and also included higher and lower concentrations (42).

Vincristine and doxorubicin can be used directly without further consideration. Cyclophosphamide, however, is a prodrug that must be metabolized by cytochrome P450 enzymes in the liver to form the active metabolite 4-OH-cyclophosphamide. For our *in vitro* study, we instead used mafosfamide, a derivative of this active compound (47). Since 4-OH-cyclophosphamide plasma levels should be transferable to mafosfamide concentrations, we used the plasma levels of 4-OH-cyclophosphamide after cyclophosphamide application as a reference (48). However, there is limited information available regarding 4-OH-cyclophosphamide plasma levels and turn-over rates from cyclophosphamide to its active form (49).

To quantify the effects of the chemotherapeutics from the R-CHOP regimen *in vitro*, we included cell lines representing the two molecular DLBCL subtypes as classified by the cell-of-origin: activated B-cell like (ABC)- and germinal-center B-cell like (GCB-type) (2). RI-1, U-2932 and TMD8 are ABC-type, and WSU-DLCL2 cells are GCB-type DLBCL. The workflow is shown in Figure 1. To assess viability and proliferation rates for the cell lines in the presence of chemotherapeutics, the number of viable cells per condition was measured at the respective time points using the CellTiter-Blue viability assay. NK cells were expanded over a period of two weeks and then incubated with the respective chemotherapeutic agents. Viability and cytotoxic capacity were quantified after 24, 48 and 72 hours of incubation. The number of viable NK cells was measured using a CASY cell counter (for 5×5 matrix incubations of NK cells, a CellTiter-Glo assay was used), and the anti-lymphoma cytotoxicity was compared in a calcein-based time-resolved killing assay (50).

**Figure 1:**
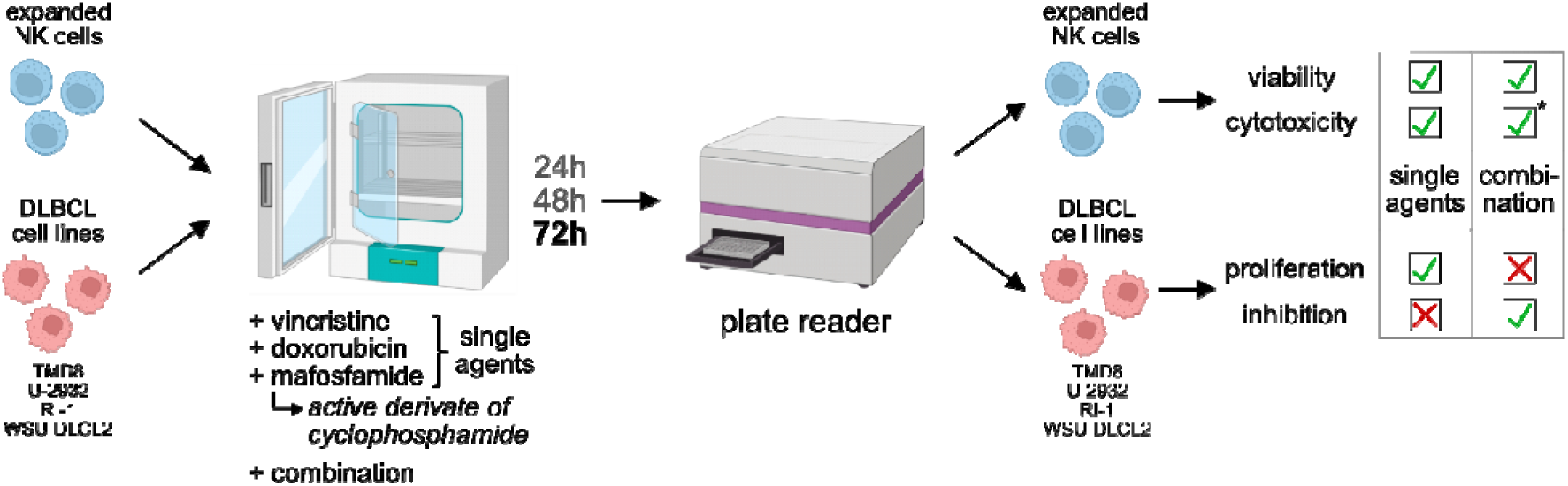
Experimental workflow. Expanded NK cells and the DLBCL cell lines (TMD8, U-2932, RI-1, WSU-DLCL2) were incubated with different concentrations of the chemotherapeutic drugs, vincristine, doxorubicin and mafosfamide, individually and in combination, for 24, 48 and 72 hours. After incubation, proliferation (for single agents) and inhibition (for combinations) were calculated for the DLBCL cell lines. The impact on NK cell viability and cytotoxicity against TMD8 cells was measured (for single agents and combinations). The asterisk marks that only certain dose combinations were further analyzed in terms of cytotoxicity.

### Effects of vincristine, doxorubicin and mafosfamide on DLBCL cell proliferation, NK cell viability and NK cell cytotoxicity against TMD8

Figure 2A-C depicts the average effects of vincristine, doxorubicin and mafosfamide on cancer cell proliferation for WSU-DLCL2, U-2932, TMD8 and RI-1 cells at 72 hours. The respective data for 24h and 48h incubation are shown in Supplementary Figure 1 and individual data points for all cell line experiments are depicted in Supplementary Figure 2.

**Figure 2:**
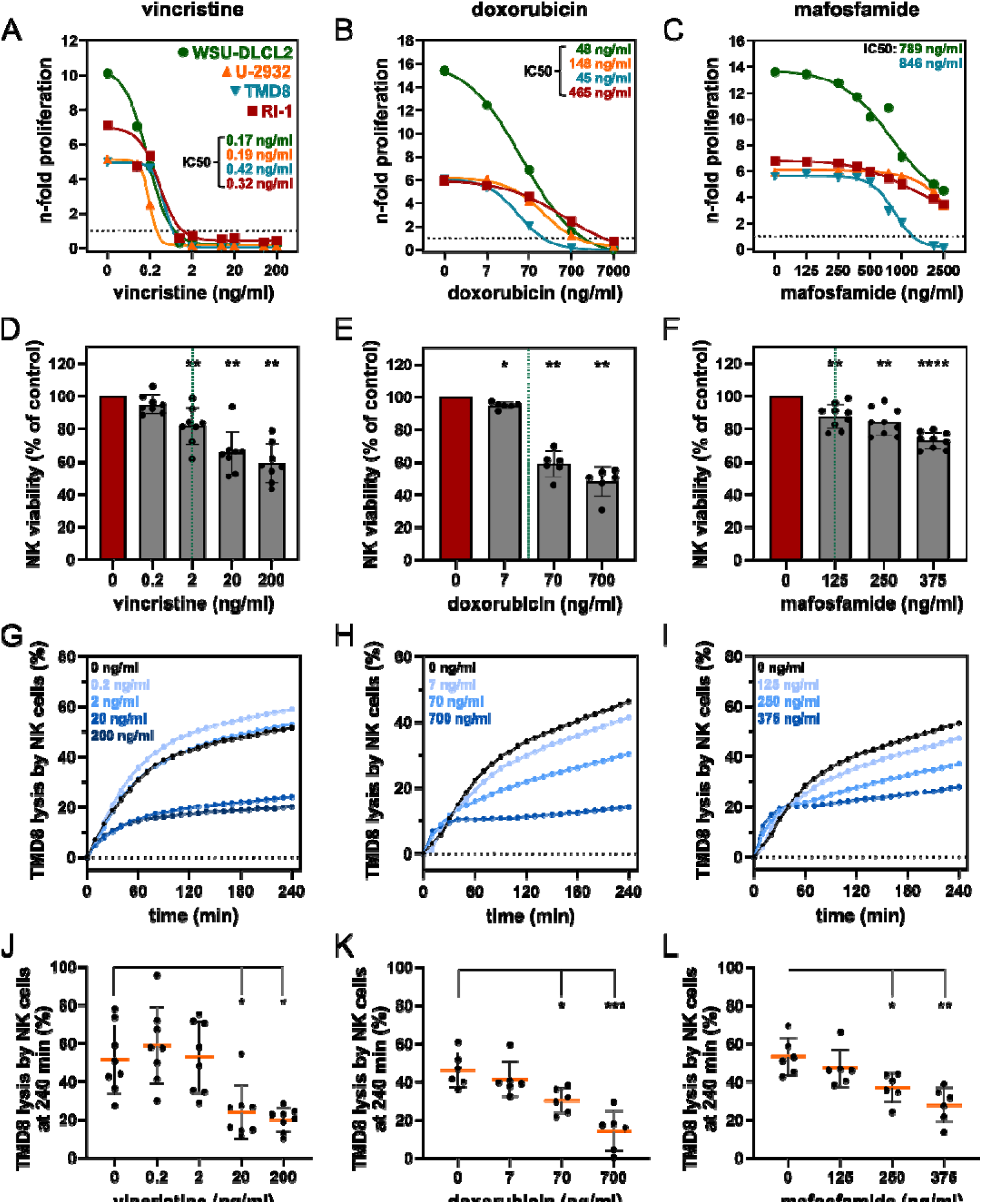
Vincristine, doxorubicin and mafosfamide reduce NK cell viability at clinical doses, but cytotoxicity only at higher doses, while effects on DLBCL cell proliferation vary widely between agents. (A-C) n-fold proliferation of the DLBCL cell lines WSU-DLCL2 (green), U-2932 (orange), TMD8 (blue) and RI-1 (red) after 72 hours of incubation with different concentrations of vincristine (A), doxorubicin (B) or mafosfamide (C). IC50 values (where calculable) are shown and are color-coded according to cell line colors. Data points represent mean values (n=4-6, see Supplementary Figure 2). (D-F) Viability of expanded NK cells after 72 hours of incubation with different concentrations of vincristine (D), doxorubicin (E) or mafosfamide (F), normalized to untreated control. Green dotted lines represent clinical plasma concentrations. Bars represent mean ± SD (n=6-9). (G-I) Mean TMD8 lysis kinetics with rituximab over 4 hours by expanded NK cells (E:T 3:1) incubated with different concentrations of vincristine (G), doxorubicin (H) or mafosfamide (I) for 72 hours, as indicated by different shades of blue in the respective graph (n=6-8). (J-L) Mean endpoint lysis ± SD of TMD8 with rituximab after 4 hours by expanded NK cells (E:T 3:1) incubated with different concentrations of vincristine (J), doxorubicin (K) or mafosfamide (L) for 72 hours (n=6-8). Asterisks highlight statistical significances. If no asterisks are shown, no statistical significance was detected.

Whereas all four cell lines proliferated over 24 to 72 hours to various degrees, cancer cell numbers were most efficiently reduced by vincristine, which could be due to both, induction of cancer cell death or inhibition of proliferation. IC50 values for vincristine after 72 hours of incubation were between 0.2 and 0.4 ng/ml for all four cancer cells lines (Figure 2A). For doxorubicin, IC50 values were rather different for the respective DLBCL cancer cell lines with values varying between 45 ng/ml and 465 ng/ml (Figure 2B). TMD8 and WSU-DLCL2 cells were more strongly affected than the two other cell lines, with IC50 values of 45 ng/ml and 48 ng/ml, respectively. Since the typical concentration of doxorubicin in patient plasma ranges from 6 ng/ml to 21 ng/ml between 1 and 25 hours after drug administration (42), we also tested doxorubicin concentrations in small increments between 0 and 50 ng/ml (Supplementary Figure 3). Doxorubicin had only minor effects on DLBCL cell proliferation within this dosage range (Supplementary Figure 3A-C). Incubation with mafosfamide, the derivative of the clinically relevant breakdown product of cyclophosphamide, had only very small effects on B cell proliferation. Mafosfamide exerted its most potent effects on WSU-DLCL2 and TMD8 cells, with IC50 values around 800 ng/ml (Figure 2C). This exceeds the clinically reported range of plasma concentrations (5.7–88.1 ng/ml).

In addition to their effects on cancer cell numbers, all substances also reduced NK cell viability in a dose-dependent manner. This effect was significant for vincristine at concentrations of 2 ng/ml or higher after 72 hours (Figure 2D). Doxorubicin incubation significantly reduced NK cell viability to <60% with concentrations of 70 ng/ml and 700 ng/ml (Figure 2E). However, even lower concentrations already affected NK cell viability (Supplementary Figure 3D-F). Mafosfamide affected NK cell viability as well with significant reductions already at doses of 125 ng/ml (Figure 2F), which is equivalent to the clinically relevant dose of cyclophosphamide.

To determine whether these effects also resulted in altered NK cell function, we quantified NK cell cytotoxicity in the presence of rituximab. For this, we incubated NK cells for 72 hours with different concentrations of the respective chemotherapeutics and subsequently analyzed their rituximab-mediated cytotoxicity against the DLBCL lymphoma cell line TMD8. This is demonstrated by the time courses of cytotoxicity over four hours (Figure 2G-I) and the quantification at 240 minutes (Figure 2J-L). As a control for NK cell functionality, we analyzed natural cytotoxicity against K562 cells without prior exposure to chemotherapeutic substances (Supplementary Figure 4). As expected, NK cells eliminated almost all K562 cells over four hours. Against TMD8 cells, NK cells were less effective but were not affected by low concentrations of 0.2 ng/ml vincristine, which is well below the standard clinical dose (Figure 2G, J). However, significant cytotoxicity impairments were observed at 20 or 200 ng/ml vincristine. As for vincristine, doxorubicin was also tested on potential inhibition of NK cell cytotoxicity. At concentrations of 70 ng/ml and higher, doxorubicin induced a significant and consistent reduction in NK cell cytotoxicity (Figure 2H, K). Similarly, incubation of NK cells with mafosfamide led to a significant reduction of cytotoxicity at concentrations of 250 ng/ml or higher (Figure 2I, L).

Similar trends and effects were observed after 24 and 48 hours of incubation for all three drugs (Supplementary Figure 1). As expected, prolonged chemotherapeutic exposure enhanced the effects on DLBCL cell proliferation, as well as NK cell viability and cytotoxicity.

### Effects of drug combinations on the number of DLBCL cells

In addition to single-drug use, we also tested the effects of chemotherapeutic combinations on DLBCL cell lines in a 5×5 matrix, leading to 125 combinations. To cover a broad range of clinically relevant doses, we chose lower concentrations than in the experiments before. We incubated DLBCL cell lines for 72 hours, measured the viability with the CellTiter-Blue assay and calculated the inhibition of DLBCL cell growth compared to untreated control. Figure 3 A-C shows two-drug combinations for TMD8 cells and Supplementary Figure 5 for RI-1, U-2932 and WSU-DLCL2 cells. Exposure with individual chemotherapeutics was also tested in this setup to compare the method to the data shown in Figure 2. Counts of all cell lines were reduced by clinical doses of vincristine and doxorubicin alone, but not mafosfamide. While doxorubicin alone reduced B-cell growth, depending on the tested cell line, by 16.3% (RI-1) to 57.3% (WSU-DLCL2) with a clinically relevant dose (19.7 ng/ml), vincristine alone inhibited growth of TMD8, U-2932 and WSU-DLCL2 completely, and RI-1 cells to 85.3% with a clinically relevant dose (2 ng/ml). Although increasing doxorubicin to 70.2 ng/ml led to a higher inhibition of all four tested cell lines, it could not reach inhibition efficiencies of vincristine alone with 2 ng/ml, in our *in vitro* setting.

**Figure 3:**
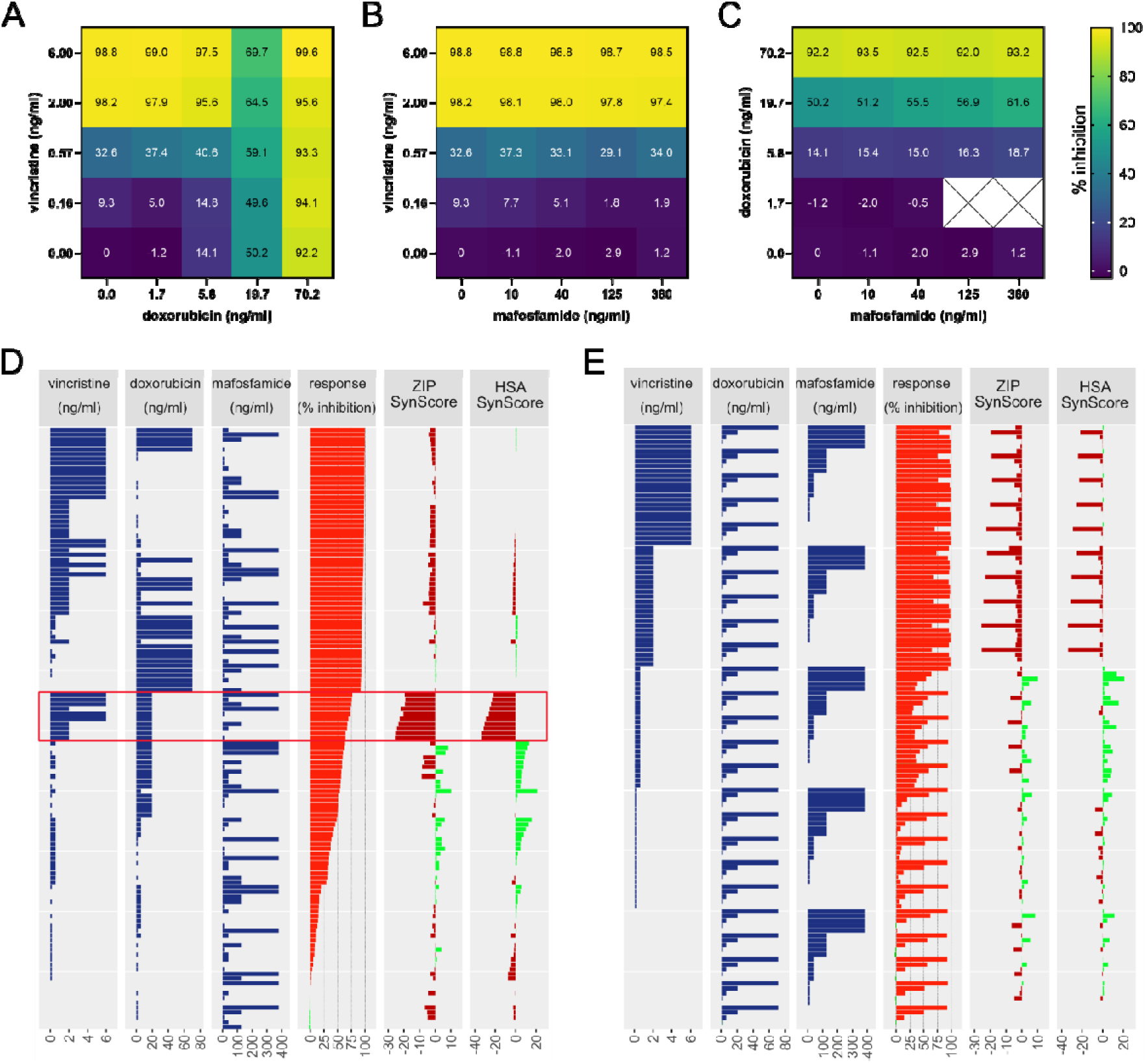
Growth inhibition of TMD8 by two- and three-drug combinations. (A-C) Heat maps for the combinations of vincristine and doxorubicin (A), vincristine and mafosfamide (B), or doxorubicin and mafosfamide (C). Color codes represent percentage of viability inhibition. TMD8 cell viability was measured after 72 hours with CellTiter-Blue (D, E) Synergy scores (ZIP and HSA) were calculated with Synergy Finder (https://synergyfinder.org/) (51) for TMD8 cells with combinatorial concentrations of vincristine, doxorubicin and mafosfamide as given in D and E. Red box in D highlights the strongest observed antagonisms. Data are sorted according to percentages of viability inhibition (D) and for vincristine concentrations (E).

The addition of vincristine and doxorubicin together increased DLBCL cell growth inhibition in a dose-dependent way, with the highest inhibition of TMD8 at 6 ng/ml vincristine and 70.2 ng/ml doxorubicin (Figure 3A). Interestingly, the combination of 2 or 6 ng/ml vincristine with 19.7 ng/ml doxorubicin was significantly less efficient in inhibiting cancer cell growth compared to vincristine alone or in combination with lower doxorubicin doses. This effect was observed for all four DLBCL cell lines and could additionally be seen at 2 or 6 ng/ml vincristine and 70.2 ng/ml doxorubicin in RI-1, U-2932 and WSU-DLCL2 cells (compare Figure 3A-C and Supplementary Figure 5). The addition of mafosfamide to vincristine did not provide an additional benefit for inhibition of TMD8 or RI-1 cells, and even further decreased inhibition of U-2932 and WSU-DLCL2 cells with increasing mafosfamide concentration, when combined with 0.57 ng/ml vincristine (Figure 3B, Supplementary Figure 5B, E, H). Combining mafosfamide and doxorubicin did not potentiate B-cell inhibition of RI-1 and U-2932 cells but slightly increased inhibition of TMD8 and WSU-DLCL2 cells, when mafosfamide was added to 5.8 or 19.7 ng/ml doxorubicin (Figure 3C, Supplementary Figure 5C, F, I). The sensitivity of all cell lines varied for different combinations, while TMD8 were most sensitive to doxorubicin and RI-1 cells were least susceptible to all three cytostatic agents.

A three-drug combination approach revealed synergistic and antagonistic effects of combinations. Synergy scores were calculated using SynergyFinder+ (Figure 3D, E, Supplementary Figure 6). Figure 3D and E show the ZIP and HSA synergy scores for TMD8 cells which either have less effect than expected (antagonism, red) or more effect than expected (synergism, green). Data are sorted according to percentages of inhibition (Figure 3D) or for vincristine concentrations (Figure 3E). Especially high negative ZIP and HSA scores were achieved for the attenuating effect when combining 2 or 6 ng/ml vincristine with 20 ng/ml doxorubicin, highlighting this antagonistic effect (red box, Figure 3D). Positive ZIP and HSA scores were also found, however, to a smaller extent. Nevertheless, they for instance support an advantageous combination of mafosfamide and doxorubicin in TMD8 and WSU-DLCL2 cells (Figure 3D, E, Supplementary Figure 6).

### Effects of drug combinations on NK cell survival and cytotoxicity

Expanded NK cells were incubated with various combinations of the three chemotherapeutics for 72 hours. In total, 125 different combinations were tested. Viability was measured using the CellTiter-Glo assay, and growths inhibition compared to untreated control was calculated (Figure 4A-C). Unexpectedly, low to medium doses of all chemotherapeutics alone had an advantageous effect on NK cell viability (vincristine: 0.1-0.6 ng/ml; doxorubicin: 2-20 ng/ml; mafosfamide: 13-125 ng/ml). For some of the low concentrations, this result differs slightly from the previous NK viability experiments, but this can be explained by different NK cell expansion growth rates and/or donor variability. Since vincristine and doxorubicin primarily target dividing cells, donor-dependent proliferation differences of NK cells can alter susceptibility to chemotherapeutics. Incubation with clinically relevant doses of doxorubicin (20 ng/ml) and mafosfamide (125 ng/ml) alone did not reduce NK cell viability in these experiments, but vincristine showed a slight decrease at a dose of 2 ng/ml. However, combinations of clinical doses of at least two substances decreased NK cell viability by approximately 30%.

**Figure 4:**
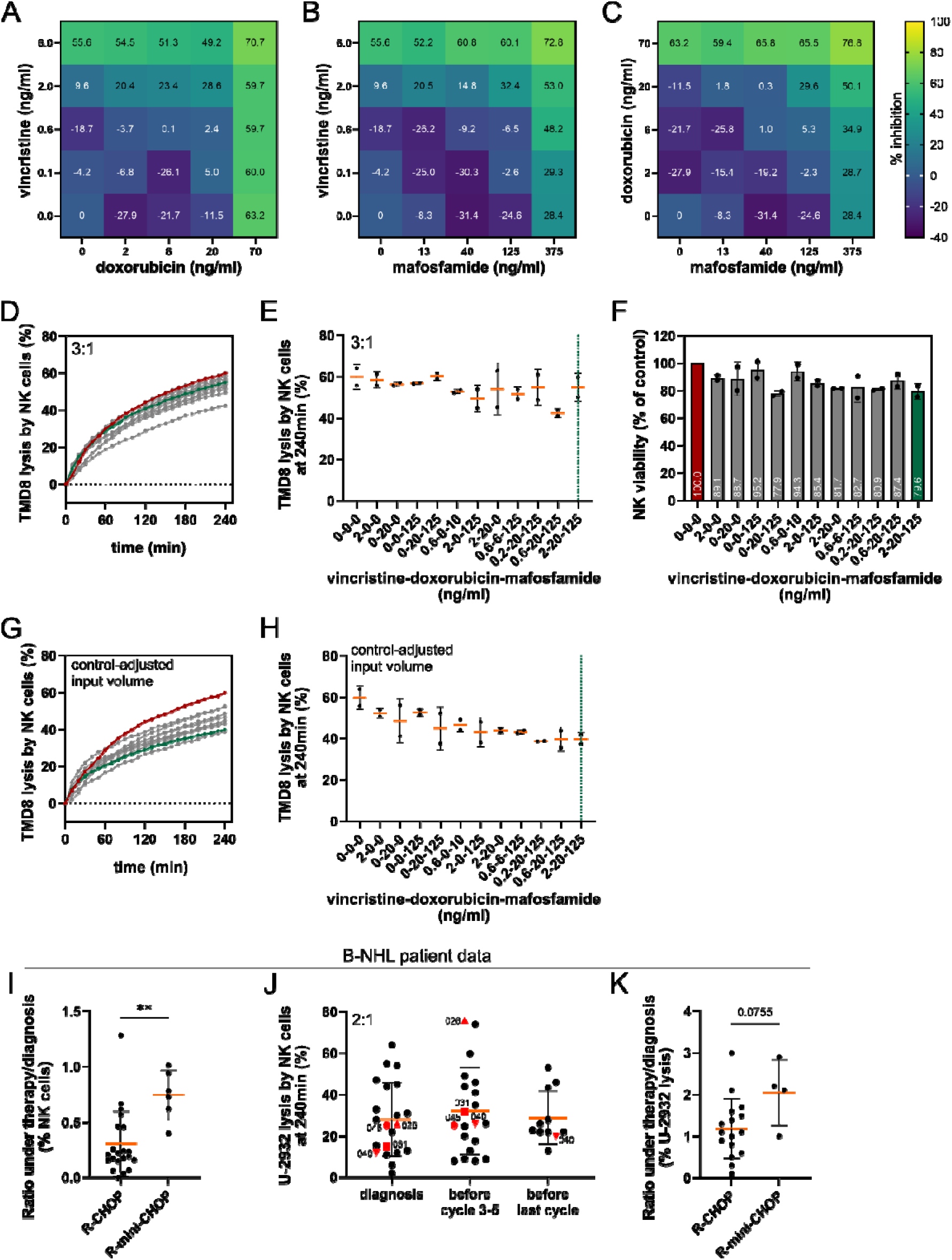
Comparison of the combinatorial applications of chemotherapeutics *in vitro* and *in vivo*. (A-C) Heatmaps showing mean inhibition of NK cell viability (%) normalized to untreated control for different concentrations and combinations of vincristine, doxorubicin and mafosfamide, after 72 hours of incubation, measured with CellTiter-Glo. Colors indicate increases in viability (less inhibition, negative values) by blue/purple, and reductions in viability (more inhibition, positive values) in green/yellow, compared to untreated control (n=2). (D, G) Mean TMD8 lysis kinetics over 4 hours with different combinations of vincristine, doxorubicin and mafosfamide concentrations incubated for 72h in expanded NK cells (n=2). Highlighted in green is the combination that equals clinically relevant doses (2-20-125), highlighted in red is the untreated control. All other concentration combinations in grey but quantified in E and H. (E, H) Mean endpoint lysis ± SD of TMD8 after 4 hours with different combinations of vincristine (V), doxorubicin (D) and mafosfamide (M) (given as V-D-M) concentrations incubated for (72h) with expanded NK cells (n=2). Green dotted lines represent clinically relevant concentrations. (D, E) Fixed E:T ratio of 3:1. (G, H) 3:1 E:T ratio was used for untreated control (0-0-0). For treatment conditions, the same volume was used as for untreated control, potentially resulting in different E:T ratios due to differences in viability. (F) Viability of expanded NK cells after 72 hours after incubation with different combinations of vincristine, doxorubicin and mafosfamide concentrations was measured with the CASY cell counter and normalized to untreated control. Highlighted in green is the combination that equals clinically relevant doses, highlighted in red is the untreated control. Bars represent mean ± SD (n=2). (I) Mean ratio ± SD of NK cell proportions (%) under therapy/diagnosis of B-NHL patients treated with R-CHOP (n=21) or R-mini-CHOP (n=6). (J) Mean endpoint lysis ± SD of U-2932 after 4 hours with primary NK cells (E:T ratio 2:1) from patients with NHL at different time points of therapy: diagnosis (n=20), before a mid-treatment R-CHOP cycle (cycle 3–5, n=19) and before last cycle (6-8, n=11). Patients receiving R-mini-CHOP are highlighted red. (K) Mean ratio ± SD of U-2932 lysis by NK cells with rituximab (%) under therapy/diagnosis of B-NHL patients treated with R-CHOP (n=16) or R-mini-CHOP (n=4).

Combinations of vincristine and doxorubicin produced a dose-dependent, additive effect on NK cell inhibition. This effect reached saturation with the highest doses of 6 ng/ml vincristine or 70 ng/ml doxorubicin (Figure 4A). Only the combination of the two highest doses further impaired NK cell viability. The combination of rather low doses of mafosfamide (13-125 ng/ml) and vincristine (0.1-0.6 ng/ml) was favorable for NK cell viability, while addition of mafosfamide to 2 or 6 ng/ml vincristine decreased NK cell viability in a dose-dependent way (Figure 4B). A similar trend was observed for the combination of mafosfamide and doxorubicin, where NK cell viability decreased with increasing doses of mafosfamide (Figure 4C). Interestingly, vincristine at the highest dose alone inhibited NK cells almost twice as much as the highest dose of mafosfamide alone. Nevertheless, the strongest NK cell inhibition was observed following combined treatment with the highest doses of doxorubicin and mafosfamide. Out of the 125 tested combinations, 12 were selected for detailed investigation of NK cell cytotoxicity based on their potential clinical relevance (compare Figure 3, Supplementary Figure 5 and 6).

Chemotherapeutics can impair NK cell cytotoxicity by reducing the number of cytotoxic NK cells, or by reducing cytotoxic capacity of the remaining living NK cells, or a combination of both. To distinguish between these effects, we prepared NK cells for two parallel killing assays. NK cells were incubated with different combinations of vincristine, doxorubicin and mafosfamide for 72 hours. In the first killing assay, viable NK cells were counted after drug incubation and used at a fixed E:T ratio of 3:1 across all conditions (Figure 4D, E; “3:1”). Untreated NK cells showed the highest rituximab-mediated cytotoxicity against TMD8 target cells (Figure 4D, red curve) whereas the drug combination of clinically relevant concentrations slightly impaired cytotoxicity (Figure 4D, green curve). However, single agents in clinically relevant concentrations only mildly impaired NK cell cytotoxicity (Figure 4E). Because viable NK cells were adjusted to a fixed 3:1 E:T ratio, the first assay specifically addressed the cytotoxic capacity of surviving NK cells independent of any chemotherapy-induced changes in NK cell counts. We next examined NK cell viability after 72-hour drug incubation to determine the extent to which chemotherapy reduced the number of available effector cells (Figure 4F). NK cell viability varied across treatment conditions. The lowest viability was observed for the clinically relevant drug combination (2 ng/ml vincristine, 20 ng/ml doxorubicin, 125 ng/ml mafosfamide, 2-20-125) (42) as well as for a condition without vincristine (0-20-125), reaching 79.6% and 77% viability, respectively. In the second “control-adjusted input volume” killing assay, we wanted to evaluate how reduced viability of NK cells affects overall target cell lysis. First, we determined the volume of viable NK cells required to achieve a 3:1 E:T ratio for the untreated control (0-0-0). For every treatment condition, we then used the same input volume of NK cells as for the control condition, resulting in condition-dependent E:T ratios due to differences in NK cell viability (Figure 4G, H; “control-adjusted input volume”). This experiment demonstrates the combined impact of chemotherapy on both NK cell viability and cytotoxic function. Consistent with the first cytotoxicity assay, untreated NK cells showed the highest rituximab-mediated cytotoxicity against TMD8 target cells (Figure 4G, red curve, H). Among single-agent treatments, vincristine (2-0-0) and mafosfamide (0-0-125) had the weakest impact on NK cell function, reducing TMD8 lysis by less than 10% in both donors. In contrast, treatment with doxorubicin alone (0-20-0) led to donor-specific responses, ranging from minimal impairment in one donor to about 20% reduction in cytotoxicity in the other (Figure 4H). Notably, despite losing 20% of viable NK cells during incubation with the clinically relevant dose combination (2-20-125) (Figure 4F; highlighted green), NK cells retained most of their cytotoxic potential, when tested at a fixed 3:1 E:T ratio (Figure 4D, E; highlighted green). However, the combined impact of reduced viability and cytotoxicity was evident in the “control-adjusted input volume” cytotoxicity assay. Here, NK cells exposed to clinically relevant doses lysed 20% fewer TMD8 cells (Figure 4G, H; highlighted green) compared to the control condition. This emphasizes that chemotherapy rather interferes with NK cell viability than with their cytotoxic function. Relevantly for the *in vivo* situation, this was mirrored in the B-NHL patient cohort we analyzed (Figure 4I-K). We compared NK cell proportions between diagnosis and under therapy, and calculated the ratio for patients receiving R-CHOP or dose-reduced R-mini-CHOP. While standard R-CHOP treatment resulted in a strong decline in circulating NK cells, R-mini-CHOP patients experienced a significantly less pronounced decline in NK cell numbers (Figure 4I). Next, we isolated primary NK cells from the peripheral blood of B-NHL patients and evaluated their cytotoxic potential against U-2932 target cells at different time points of R-CHOP therapy. At time of diagnosis, no chemotherapeutics were given yet, in contrast to mid-therapy (before cycle 3-5) and before the last cycle (before cycle 6-8). In agreement with our *in vitro* data (Figure 4E), R-CHOP application did not alter the cytotoxic capacity of circulating patient NK cells, at any of the investigated time points during chemotherapy compared to the time of diagnosis (Figure 4J). As expected, the cytotoxic activity against U-2932 cells was also preserved during R-mini-CHOP treatment but notably showed a trend toward increased cytotoxicity over the course of therapy (Figure 4K).

Together, these findings demonstrate that clinically relevant R-CHOP-associated cytostatic exposure primarily affects NK cell viability rather than their intrinsic cytotoxic function. Although reduced NK cell numbers translated into diminished overall killing capacity in the “control-adjusted input volume” experiment, viable NK cells largely retained their cytolytic potential both *in vitro* and throughout therapy in B-NHL patients (Figure 4D, E, G-I). These results suggest that impaired NK cell-mediated anti-lymphoma activity during treatment is mainly driven by quantitative rather than qualitative defects in the NK cell compartment.

### Balancing efficacy and toxicity in R-CHOP: A mathematical model

Achieving the right balance between eliminating target cells efficiently and ensuring optimal survival of effector cells is one of the central challenges of R-CHOP therapy. While maximizing cytotoxic activity against malignant B-cells is essential for therapeutic efficacy, excessive treatment intensity simultaneously impairs viability of immune effector cells that are required for sustained anti-tumor responses, especially when combined with immunotherapeutic approaches. This trade-off adds considerable complexity to the system, and thus also to the treatment of patients. To address this challenge, we developed a mathematical optimization model based on pareto-optimality and integrated our *in vitro* data (Figure 5). The model describes the balance between cancer and effector cell survival by identifying treatment configurations that optimize both objectives simultaneously. In Figure 5A-C, cubic polynomial surfaces are shown for combinations of two chemotherapeutics, respectively. The model suggests optimal concentrations of vincristine (2.1 ng/ml), doxorubicin (0 ng/ml), and mafosfamide (21.7 ng/ml) to eliminate nearly all target cells and preserve as many NK cells as possible.

**Figure 5:**
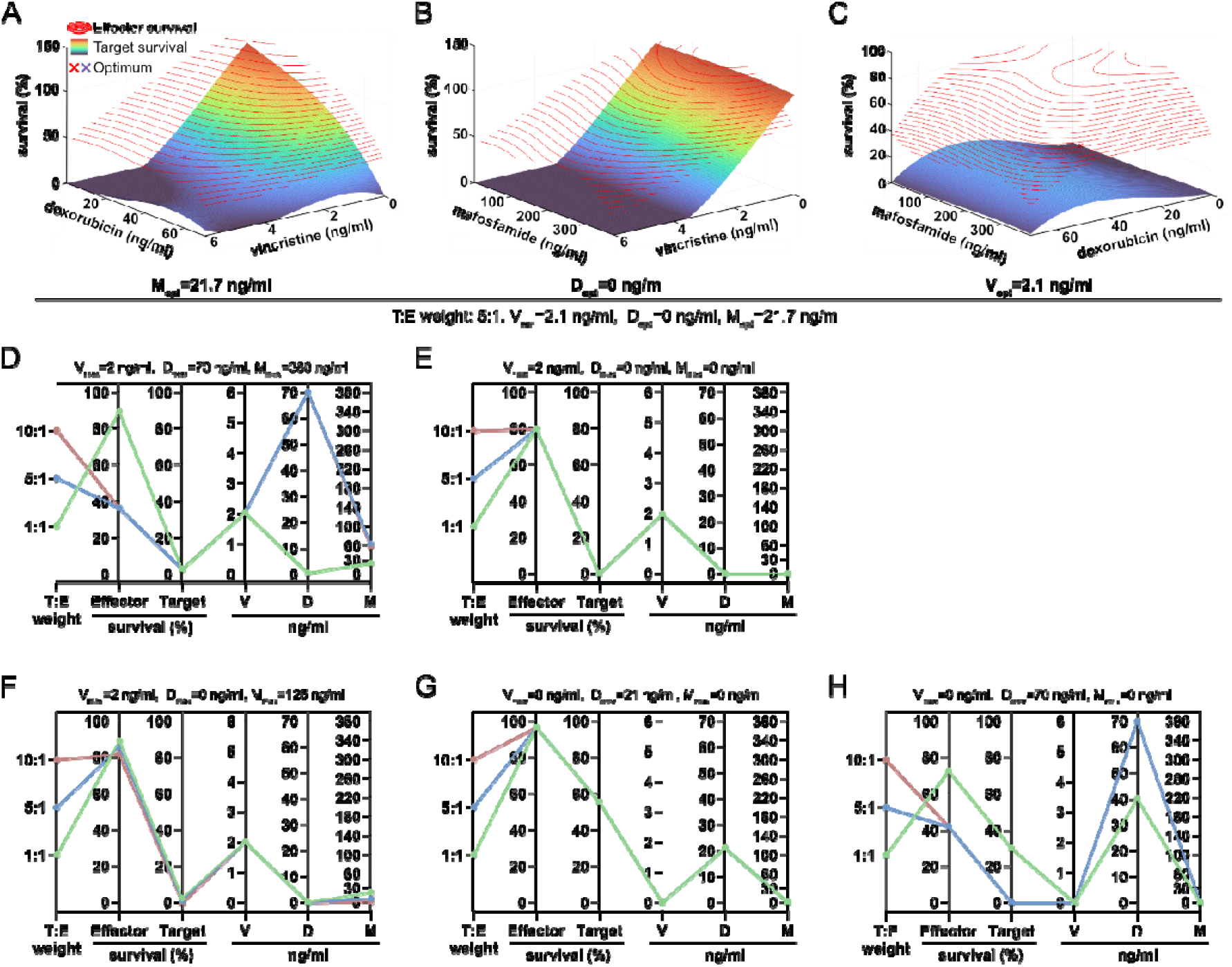
Optimization of target cell elimination and effector survival rates during R-CHOP. Vincristine (V), doxorubicin (D) and mafosfamide (M) doses are plotted as ng/ml. (A-C) Rainbow-colored, cubic polynomial surfaces describe target survival (%). The red lines on top describe NK cell survival rates (%). The polynomial surfaces were modeled for T:E weight (target:effector weight) 5:1, V_opt_ = 2.1 ng/ml, D_opt_ = 0 ng/ml M_opt_ = 21.7 ng/ml. Each 3D surface includes the dose range of two chemotherapeutics and sets the third dose on the respective computed optimum. A red or purple cross (behind/in front of the polynomial surface) marks the optimal doses for the plotted agents, as calculated by our model. (D-H) Parallel coordinate plots are based on chosen limitations, as indicated on top of each panel (V_max_=X; D_max_=Y; M_max_=Z). Plots include optimal target elimination and NK cell survival rates, as well as potential dose ranges for vincristine (V), doxorubicin (D) and mafosfamide (M). Depending on T:E weight (1:1=green; 5:1=blue; 10:1=red), different paths show the respective pareto-optimal solution (D-H).

Pareto-optimal solutions for certain target-to-effector survival rate weights (T:E) and limitations are highlighted in parallel coordinate plots (Figure 5D–H). To achieve this, we incorporated the full concentration range of the concentrations of the chemotherapeutics, which had been tested experimentally. When target cell death and NK survival were given equal weighting (T:E weight 1:1), the model identified a solution that was predominantly characterized by vincristine exposure. Doxorubicin was largely excluded in order to preserve effector survival, and mafosfamide contributed only minimally (Figure 5D, T:E weight 1:1). Prioritizing target death over effector survival (T:E weight of 5:1 or 10:1), the model suggests higher doxorubicin and mafosfamide concentrations together with 2 ng/ml vincristine at the expense of NK cell survival. Restricting the model to vincristine alone, capped at clinically relevant concentrations (2 ng/ml), still resulted in highly effective target cell elimination while only mildly affecting effector cell survival (Figure 5E). Based on the observed antagonism between vincristine and doxorubicin, doxorubicin was excluded from the model, yet an efficient combination of vincristine and mafosfamide still yielded very good results within clinically constrained dosage ranges (Figure 5F). Additional models were then generated in which vincristine and mafosfamide were excluded entirely, leaving doxorubicin as the sole active compound (Figure 5G, H). Under clinically capped conditions (Figure 5G), the predicted efficacy remained limited. However, extending the allowed concentration range caused the model to select substantially higher doxorubicin doses to achieve improved target cell elimination. Again, this came at the expense of NK cell survival (Figure 5H).

## Discussion

The combination of chemotherapy and immunotherapy has become the standard treatment for various forms of cancer. However, the CHOP regimen was developed long before immunotherapy became a treatment option. Thus, it is clear that the CHOP regimen was never intended to preserve the viability or functionality of immune cells, but rather to reduce tumor mass. The introduction of rituximab to the CHOP regimen (R-CHOP) efficiently improved the outcome of patients with DLBCL compared to CHOP alone (22). This made NK cell survival and functionality a fundamental requirement for rituximab therapy success. In principle, different combinations of modalities such as chemotherapy, radiation or immune therapies, as well as different concentrations, dosages, or scheduling can be optimized to improve immune cell function. However, the possibility of testing many different combinations in clinical studies is, understandably, rather limited. Thus, detailed independent preclinical data are needed to inform clinical studies. R-CHOP has been the standard first-line therapy for non-Hodgkin lymphomas such as DLBCL (22–26), but a growing number of promising new immunotherapeutic approaches are emerging. For instance, R-Pola-Glo combines rituximab with the bispecific antibody glofitamab (Glo; CD20xCD3) and the antibody-drug conjugate polatuzumab vedotin (Pola; anti-CD79B). However, these are not yet approved for use as first-line treatment. However, this “chemo-light” regimen produced even better results in a phase II clinical study involving elderly and frail patients than other conventional therapies would in this risk group (52, 53).

We investigated the effects of R-CHOP therapy on NK cells and four DLBCL cell lines *in vitro*. This treatment regimen combines rituximab with vincristine, doxorubicin, and cyclophosphamide, administered every 2-3 weeks at standard doses used in first-line non-Hodgkin lymphoma therapy. Reported plasma concentrations 1 to 25 hours after administration range from 0.9-2.6 ng/ml for vincristine, 6-21 ng/ml for doxorubicin, and 5.7-88.1 ng/ml for 4-OH-cyclophosphamide (48). Vincristine, doxorubicin, and mafosfamide (a derivative of the biologically active form of cyclophosphamide) reduced numbers of cells in all DLBCL cell lines to varying degrees at concentrations achieved during R-CHOP therapy. Vincristine was the most effective agent, with IC50 values below 0.5 ng/ml and near-complete eradication of all cell lines at 2 ng/ml after 72 hours, matching reported plasma concentrations 1 to 25 hours after administration (41). Doxorubicin was less effective, showing IC50 values of 45 ng/ml in TMD8 and 48 ng/ml in WSU-DLCL2 cells, but substantially higher values in the other cell lines after 72 hours (up to 465 ng/ml for RI-1). As TMD8 originated from an untreated patient (54), whereas the other cell lines were derived from pretreated patients (55–57), acquired resistance may explain these differences, although intrinsic susceptibility cannot be excluded. For mafosfamide, IC50 values could only be determined for two DLBCL cell lines in the tested concentration range and were around 800 ng/ml for mafosfamide, which is rather high considering the reported plasma levels of 4-OH-cyclophosphamide (5.7-88.1 ng/ml; 1 hour after administration) (48).

As expected, all three drugs reduced NK cell viability in a dose-dependent manner, sometimes even at the lowest tested concentration. NK cytotoxicity against TMD8 cells in the presence of rituximab was also compromised, most notably by mafosfamide and, at higher concentrations, by vincristine and doxorubicin. Even at the lowest doxorubicin concentration (7 ng/ml), a slight inhibitory trend was observed whereas low vincristine concentrations (0.2 ng/ml) had no negative effect.

Our combinatorial experiments covered the reported plasma concentrations (see above) and demonstrated that clinically relevant dose of vincristine (2 ng/ml) alone was sufficient to inhibit DLBCL growth *in vitro*, without further enhancement by doxorubicin and only a very slight impact of mafosfamide, as also demonstrated by the mathematical model. Instead, antagonistic effects, particularly in combination with 20 ng/ml doxorubicin, were observed across all four cell lines. This antagonism between vincristine and doxorubicin has been reported previously by Erhardt *et al.* and depends on treatment timing and mechanistic interactions (40). Overall, our data revealed little evidence for synergistic effects among the tested R-CHOP components in DLBCL cell lines.

The NK cell experiments with single agents and their combinations demonstrated that clinically relevant doses impair NK cell viability and, to a certain extent, cytotoxic function. While mafosfamide alone reduced NK cell cytotoxicity, vincristine and doxorubicin alone did not show this effect when applied as single agents *in vitro*. Combinations of clinical doses did not significantly reduce cell cytotoxicity when compared to control. In contrast, single agents and combinations always reduced NK cell viability at clinical concentrations. This is consistent with reduced viable NK cell counts measured in our B-NHL cohort throughout the course of treatment (40) and previous reports showing a decline in NK cell numbers during therapy (58). Combining two or more agents at clinical doses led to a more pronounced reduction in NK cell viability compared to single agent treatment. This was also illustrated by the mathematical model, in which a high T:E weight favored a combination of all three agents but at the expense of NK cell survival (Figure 5D). In contrast, a T:E weight of 1:1 suggested the omission of doxorubicin and the use of a combination of 2 ng/ml of vincristine and low doses of mafosfamide, preserving over 80% of NK cells. Lower drug concentrations largely preserved NK cell cytotoxicity under single-agent conditions, and viability across both, single-agent and combinatorial treatments. However, these low doses were insufficient to effectively inhibit tumor cell growth *in vitro*. Given that viable NK cells largely retained their cytotoxic potential both after *in vitro* drug exposure and throughout the course of immunochemotherapy in B-NHL patients, our data indicate that clinically relevant R-CHOP doses primarily compromise NK cell viability rather than the intrinsic cytotoxic function of surviving NK cells. These findings were further supported by our observation that the dose-reduced R-mini-CHOP regimen preserved NK cell numbers more effectively throughout treatment than standard-dose R-CHOP. Notably, patients receiving R-mini-CHOP exhibited a trend toward enhanced NK cell cytotoxicity during therapy, while standard dose R-CHOP neither reduced nor increased NK cell cytotoxicity.

Considering only the *in vitro* data, omitting doxorubicin and/or cyclophosphamide and thereby limiting treatment to rituximab and vincristine could be an option. Although there is little data comparing monotherapy with combined therapy involving vincristine, doxorubicin and cyclophosphamide, clinical experience and several other arguments clearly speak against monotherapy with vincristine. In a rapidly proliferating and heterogeneous disease such as DLBCL, combinatorial treatment strategies are essential to prevent diverse escape mechanisms. Early foundational work by Luria and Delbrück (59) demonstrated that bacteria resistance arises spontaneously rather than in direct response to selective pressure. Building on this concept, Law extended these findings to chemotherapy, showing that resistant tumor cell clones frequently pre-exist prior to treatment (60). Gatenby *et al*., postulated that sensitive and resistant cells compete for resources, meaning that survival of sensitive cells suppresses proliferation of resistant clones (61). Consequently, monotherapy imposes a one-dimensional selection pressure, favouring the outgrowth of resistant clones. In contrast, combination therapy such as R-CHOP mitigates this risk by targeting multiple pathways simultaneously (62, 63).

Taken together, these findings emphasize the difficulty of balancing effective tumor control while preserving immune effector function. Although our *in vitro* data suggest that reduced-intensity regimens may better preserve NK cell viability and cytotoxic function, such approaches are unlikely to provide durable disease control in a clinically relevant setting. However, the success of chemo-free or chemo-light regimens, such as R-Pola-Glo, suggests that minimizing immunotoxicity does not necessarily compromise efficacy. Taking all considerations into account, we propose studying a modified treatment schedule, where doxorubicin application is postponed to a later time point, while the remaining R-CHOP components are administered upfront (as modelled in Figure 5F-H). This approach is based on the rationale that early preservation of NK cell viability and function may enhance rituximab-mediated ADCC during the initial phase of treatment, while maintaining overall anti-tumor efficacy. Potentially, this regimen could also avoid antagonistic effects between vincristine and doxorubicin, as reported by Erhardt *et al*., thus promoting tumor control (64). Similarly, alternative strategies such as antibody-drug conjugates, nanoformulations and liposomal drug delivery systems may help to maintain anti-tumor efficacy while limiting off-target effects on immune cells.

Within this evolving therapeutic landscape, optimizing the interplay between cytotoxic agents and the immune system may represent a key step toward improving outcomes in DLBCL. Although the proposed schedule is hypothetical and requires rigorous clinical validation, it shows how conventional regimens could be improved to make better use of immunological mechanisms and support the effectiveness of modern immunotherapy-based treatments.

## Materials and Methods

### Ethical approval

Experiments with mononuclear blood cells and patient samples are approved by the local ethics committee (reference 84/15 and 33/18). The local blood bank of the Institute of Clinical Hemostaseology and Transfusion Medicine at Saarland University Medical Centre provided leucocyte reduction system (LRS) chambers, a byproduct of platelet collection from healthy blood donors. All blood donors and patients provided informed, written consent to use their blood for research purposes.

### Cell Lines

RI-1 (ACC 585, DSMZ), WSU-DLCL2 (ACC 575, DSMZ), U-2932 (ACC 633), K562 (CCl-243) and TMD8 cells (kindly provided by Lorenz Trümper, Göttingen) were used. K562-LucpC clone G4 and TMD8-LucpC clone F7 were generated using lentiviral transduction as previously described (40, 65), and single-cell clones were generated. All cell lines were cultured at 37°C and 5% CO_2_ in RPMI-1640 (ThermoFisher Scientific, Waltham, USA), supplemented with 10% FBS and 1% penicillin/streptomycin (P/S; Sigma-Aldrich, St. Louis, USA).

### Preparation and maintenance of NK cell expansions from blood donors

Human peripheral blood mononuclear cells (PBMC) were isolated from leukocyte reduction system chambers (LRS) using density gradient centrifugation (leukocyte separation medium LSM 1077, PromoCell, Heidelberg, Germany), as described before (66).

For the NK cell expansion, irradiated K562 feeder cells (90 Gy) and PBMCs were incubated together in a 2:1 ratio in RPMI-1640, supplemented with 10% FBS, 1% P/S and 200 U/ml IL□2 (Miltenyi Biotec, Bergisch Gladbach, Germany). Medium was changed upon pH indication, and diluted to 2.5×10^6^ cells, if necessary. After two weeks of expansion, NK cells were ready to be used.

### Incubation of expanded NK cells with chemotherapeutics

For the respective time points (24h, 48h, 72h), NK cells were seeded out with a density of 1.2×10^6^ cells/cm^2^ at a concentration of 2.4×10^6^ cells/ml, and incubated in RPMI-1640, supplemented with 10% FBS, 1% P/S and 200 U/ml IL□2 at 37°C and 5% CO_2_. The chemotherapeutic drugs vincristine (local pharmacy, Universitätsklinikum des Saarlandes, Homburg, Germany), doxorubicin (local pharmacy, Universitätsklinikum des Saarlandes, Homburg, Germany) and mafosfamide sodium salt (Santa Cruz Biotechnology, Dallas, USA) were added to the different conditions and incubated for 24, 48 and 72 hours. To exclude dimethyl sulfoxide (DMSO)-dependent effects, all conditions contained the same amounts of DMSO. For every time point, the viability of expanded NK cells was measured, using a CASY cell counter (OLS, Bremen, Germany).

### Plate printing for combinatorial incubations of chemotherapeutics

Cell suspensions of DLBCL cell lines or expanded NK cells (200 µl per well) were added to 96-well plates (Corning, Corning, USA). TMD8, U-2932 and RI-1 were seeded out with 0.05×10^6^ cells/well, WSU-DLCL2 cells with 0.03×10^6^ cells/well and NK cells with 0.015×10^6^ cells/well. Three compounds were then printed on the plates using the D300e Digital Dispenser (Tecan, Männedorf, Switzerland), following the protocol detailed by Pauck *et al*. (67). Vincristine and doxorubicin were purchased from MedChemExpress (Monmouth Junction, USA) as 10 mM stock solutions. Mafosfamide was used as previously described. All compounds were solved in 100% DMSO. Stock solutions were aliquoted and diluted with DMSO to achieve optimal dispensing volumes for 96-well plate assays. Final stock concentrations for dispensing with the D300e Digital Dispenser were 0.81 mg/ml (vincristine), 5.8 mg/ml (doxorubicin), and 10 mg/ml (mafosfamide). Different compounds were printed in each well determined by a 5×5 matrix. The final DMSO concentration was 0.005%.

### Non-Hodgkin lymphoma patient cohort

All patients included in this study were diagnosed with aggressive B-cell non-Hodgkin lymphoma and received 6-8 cycles of R-CHOP (20 patients), R-CHOEP (5 patients), R-mini-CHOP (6 patients) or R-pola-CHP (1 patient). The cohort compromised 32 patients recruited at the Department of Internal medicine I (Homburg, Saarland University) between August 2019 and February 2025. PBMCs were isolated (see section above) from these patients at different time points during therapy. The proportion of NK cells within the PBMC population was determined by flow cytometry. In addition, NK cells were isolated by removing monocytes, granulocytes and T cells using human CD14, CD15 and CD3 microbeads (Miltenyi Biotec, Bergisch Gladbach, Germany), followed by the isolation of CD56-positive cells using human CD56 microbeads (Miltenyi Biotec, Bergisch Gladbach, Germany). Following overnight culture in AIMV/10% FBS, ADCC capacity of NK cells was determined using a real-time killing assay (see section below) at an effector-to-target cell ratio of 2:1. U-2932 cells were used as the target cell line. Patients in this study are predominantly part of the recently published cohort by Zöphel *et al.* (40).

### CellTiter-Blue viability assay with DLBCL cell lines

TMD8, U-2932 and RI-1 cells were seeded out with 0.05×10^6^ cells/well, WSU-DLCL2 with 0.03×10^6^ cells/well, on 96-well plates (Corning, Corning, USA) and incubated with the respective chemotherapeutics for 24, 48 and 72 hours. Three hours prior to measuring, 20 µl CellTiter-Blue reagent (Promega, Madison, USA) was added to the plates. Plates were measured in a Tecan infinite M200pro plate reader, using i-control 1.9 software (Tecan, Männedorf, Switzerland).

### CellTiter-Glo viability assay with expanded NK cells

Expanded NK cells were seeded out with 0.015×10^6^ cells/well in an opaque 384-well plate (Corning, Corning, USA) and incubated with the respective chemotherapeutics for 72 hours. Prior to measuring, 30 µl of CellTiter Glo reagent (Promega, Madison, USA), 1:1 diluted with sterile PBS, was added to each well using the Multidrop Combi (ThermoFisher Scientific, Waltham, USA). Measurements were performed in a Tecan Spark Cyto plate reader (Tecan, Männedorf, Switzerland) with an integration time of 500 ms per well.

### Calcein-based real-time cytotoxicity assay

NK cell cytotoxicity was analyzed using a calcein-based real-time assay as previously described (50). For ADCC analysis, TMD8 were used and as control for NK cell killing capacity, K562 were used. Therefore, target cells (TMD8-LucpC clone F7 and K562-LucpC clone G4) were stained with calcein-AM (500 nM, ThermoFisher Scientific, Waltham, USA) for 15 minutes at room temperature in the dark under gentle agitation. Cells were then washed with AIMV medium without FCS (ThermoFisher Scientific, Waltham, USA) and seeded at 0.025×10^6^ cells per well in 200 µl AIMV (for ADCC 1 µg/ml rituximab (10 mg/ml stock solution) was added) into a black 96-well plate with clear bottom (Corning, Corning, USA). Plates were incubated for 30 minutes at 37°C and 5% CO_2_ to allow cell settling. Expanded NK cells, previously incubated for 24, 48 or 72 hours with different concentrations of chemotherapeutic agents, as described above, were used as effector cells. NK cells were added in an effector to target (E:T) ratio of 3:1 by carefully dispensing 0.075×10^6^ viable NK cells in 50 µl AIMV per well without disturbing the target cell layer. Calcein fluorescence was measured every ten minutes over a period of four hours using a CLARIOstar microplate reader (bottom reading mode, 37°C, 5% CO_2_, BMG Labtech, Ortenberg, Germany). NK cell cytotoxicity was determined by measuring the target cell lysis and calculated as follows:

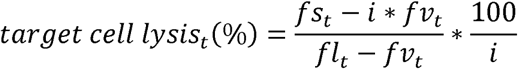

with

*fs_t_* = fluorescence of the sample at time *t*

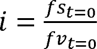 is the index at time 0

*fv_t_* = fluorescence of the viable control (target cells + 50 µl AIMV) at time *t*

*fl_t_* = fluorescence of the lysis control (target cells + 50 µl AIMV + 0,8% Tergitol 15-S-9 (Carl Roth, Karlsruhe, Germany))

### Mathematical optimization model

The objective of the optimization is to minimize target cell survival, while simultaneously maximizing NK cell survival. This poses a bi-objective optimization problem. The sought-after solution consists of Pareto-optimal combinations, meaning that it is not possible to improve one condition without worsening the other. This represents the best possible compromise. However, a minimal survival rate of target cells is more desirable than preserving the maximum number of NK cells. To achieve this, weighing factors are introduced to put more emphasis on one or the other of these objectives. Since cell viability cannot fall below 0%, it must be ensured that calculated survival rates stay within biologically meaningful boundaries. In addition, the maximum dosage of a compound delivered to a patient must be limited, due to patient safety considerations. The model therefore must support the possibility of defining individual maximum doses for each compound. Experimental conditions are treated as a vector of the form ***c*** = (*c_V_, c_D_, c_M_*), representing concentrations of vincristine (V), doxorubicin (D) and mafosfamide (M). Responses *y_T_*(***c***) and *y_NK_*(***c***) are measured as percentages of target and NK cell survival, respectively. Dose responses are modeled as a four-dimensional cubic polynomial surface of the form

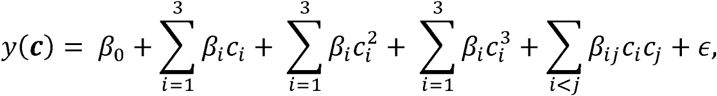

where *c_i_* are the log-transformed concentrations, *β_i_* are the regression coefficients and is the residual error. The log transform is used to make the dose-response relationship approximately linear to account for non-linear effects in drug response and large differences in dose-concentrations. To obtain a single optimal solution, responses are transformed into desirability functions. To maximize NK survival:

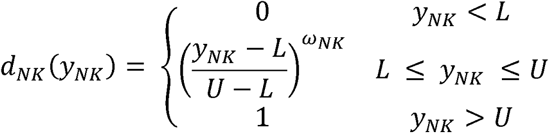

And to minimize target cell survival:

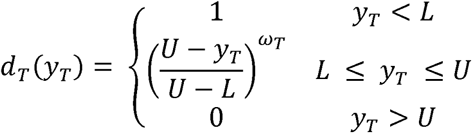

*L* and *U* are lower and upper boundaries, respectively, defining the limits of acceptable survival rates of target and NK cells, and *ω_NK_* and *ω_T_* are weights (e.g. if low cancer survival is more important set *ω_T_* > 1 and *ω_NK_* = 1). In addition, it is desirable to administer low overall doses of cytotoxic and immunosuppressive compounds. We therefore add a soft constraint that penalizes higher overall dosages of compounds *c_i_*:

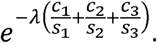

Here, *λ* is the overall scaling factor of dose penalty and are compound-dependent individual scaling factors, set to the afore-mentioned dose limits. This ensures biologically meaningful scaling of dosages by simultaneously avoiding the dominance of coupons requiring higher concentrations. Overall, the desirability function has the following form:

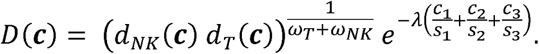

The optimal drug combination is then obtained by

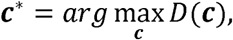

which allows to prioritize tumor growth inhibition while preserving NK viability by adjusting desirability weights. Input data for the model were experimental data of the 5×5 matrix measured for NK cells and TMD8 cell line.

### Chemicals and reagents

The chemotherapeutics vincristine and doxorubicin were purchased from the local pharmacy (Universitätsklinikum des Saarlandes, Homburg, Germany) in aqueous solutions of 1 mg/ml and 2 mg/ml, respectively. The anti-CD20 antibody rituximab (as biosimilar Rixathon (Hexal)) was purchased from the local pharmacy (Universitätsklinikum des Saarlandes, Homburg, Germany) in aqueous solutions of 10 mg/ml. Additionally, stocks in DMSO were purchased for vincristine and doxorubicin from MedChemExpress (Monmouth Junction, USA) as 10 mM stock solutions. Mafosfamide sodium salt (Santa Cruz Biotechnology, Dallas, USA) was diluted to 10 mg/ml stock solutions with DMSO. For printing of combinatorial incubations of cytostatic compounds, the DMSO-solved reagents were used. For all other experiments, aqueous solutions of vincristine and doxorubicin were used.

### Data and statistical analysis

Data are presented as mean values ± standard deviation (SD). Dots represent individual data points (except for CellTiter-Blue assays, where they represent the mean). N indicates the number of experiments or different PBMC donors, as specified. Statistical analyses were performed using GraphPad Prism (version 11.0.0, GraphPad Software Inc., Boston, USA).

Differences between treatment conditions were analyzed using one-way repeated-measures analysis of variance (RM-ANOVA) if the assumptions for parametric testing were met. Otherwise, the non-parametric Friedman test was applied. In all analyses, measurements from different treatment conditions were obtained from the same donors and were therefore treated as paired. When individual treatment conditions were missing, the mixed-effects model was used to calculate significant differences, instead of RM-ANOVA.

When a significant overall effect was detected, post hoc multiple comparisons were performed using Dunnett’s test for parametric, or Dunńs test, for nonparametric data, to compare treatment conditions to untreated control. Normality was assessed using Shapiro-Wilk Test and by visual control of Q-Q plots. Significance levels are: *p ≤0.05, **p ≤0.01, ***p ≤0.001, ****p ≤0.0001.

Synergy analyzes were performed using SynergyFinder+ (51) applying ZIP (68) and HSA synergy scoring. Microsoft Excel (version 365, Microsoft Corporation, Redmond, USA) was used for data preprocessing. Figure 1 was created in BioRender (created in BioRender. Jansky, J. (2026), https://BioRender.com/sinwm4a. Figures were finalized using Affinity Designer (version 2.4, Serif Ltd, Nottingham, UK).

## Supporting information

Jansky_Supplementary

## Author contributions

JJ, LT, ECS and MH designed the study. JJ, HB and LD performed all experiments if not otherwise mentioned. FW, DP and MR performed plate printing for combinatorial incubations of chemotherapeutics. HE provided blood samples. JJ, HB and ECS analyzed all experiments. SZ, supported by NK, prepared isolations and experiments with primary B-NHL patient-derived NK cells. GS performed mathematical modelling and simulations. JJ, HB, ECS, MB and MH co-wrote the paper. All authors critically discussed data analysis, and discussed and revised the manuscript.

## Acknowledgements

We thank the Institute of Clinical Hemostaseology and Transfusion Medicine, Saarland University and University Medical Center, Homburg, Germany for providing LRS chambers. We gratefully acknowledge Gertrud Schäfer, Kathleen Seelert, Kathrin Förderer, Cora Hoxha and Sandra Janku for cell preparation and help with solutions, ordering, and maintaining a well-functioning lab. We acknowledge Carsten Kummerow and Lennart Tegethoff for their scientific discussions of the project.

## References

1. Zöphel S, Schwär G, Nazarieh M, Konetzki V, Hoxha C, Meese E, et al. Identification of molecular candidates which regulate calcium-dependent CD8+ T-cell cytotoxicity. bioRxiv. 2021:2020.12.22.423945. doi: 10.1101/2020.12.22.423945

2. Miao Y, Medeiros LJ, Li Y, Li J, Young KH. Genetic alterations and their clinical implications in DLBCL. Nat Rev Clin Oncol. 2019;16(10):634–52. doi: 10.1038/s41571-019-0225-1

3. Stegemann M, Denker S, Schmitt CA. DLBCL 1L-What to Expect beyond R-CHOP? Cancers (Basel). 2022;14(6). doi: 10.3390/cancers14061453

4. Adair FE, Bagg HJ. Experimental and Clinical Studies on the Treatment of Cancer by Dichlorethylsulphide (Mustard Gas. Ann Surg. 1931;93(1):190–9. doi: 10.1097/00000658-193101000-00026

5. Dy GK, Adjei AA. Systemic cancer therapy: evolution over the last 60 years. Cancer. 2008;113(7 Suppl):1857–87. doi: 10.1002/cncr.23651

6. Matthias JQ, Misiewicz JJ, Scott RB. Cyclophosphamide in Hodgkin’s disease and related disorders. Br Med J. 1960;2(5216):1837–40. doi: 10.1136/bmj.2.5216.1837

7. Bohannon RA, Miller DG, Diamond HD. Vincristine in the treatment of lymphomas and leukemias. Cancer Res. 1963;23:613–21. doi:

8. Gottlieb JA, Gutterman JU, McCredie KB, Rodriguez V, Frei E, 3rd. Chemotherapy of malignant lymphoma with adriamycin. Cancer Res. 1973;33(11):3024–8. doi:

9. Luce JK, Gamble JF, Wilson HE, Monto RW, Isaacs BL, Palmer RL, et al. Combined cyclophosphamide vincristine, and prednisone therapy of malignant lymphoma. Cancer. 1971;28(2):306–17. doi: 10.1002/1097-0142(197108)28:2<306::aid-cncr2820280208>3.0.co;2-n

10. Skarin AT, Pinkus GS, Myerowitz RL, Bishop YM, Moloney WC. Combination chemotherapy of advanced lymphocytic lymphoma. Importance of histologic classification in evaluating response. Cancer. 1974;34(4):1023–9. doi: 10.1002/1097-0142(197410)34:4<1023::aid-cncr2820340410>3.0.co;2-7

11. Schein PS, DeVita VT, Jr., Hubbard S, Chabner BA, Canellos GP, Berard C, et al. Bleomycin, adriamycin, cyclophosphamide, vincristine, and prednisone (BACOP) combination chemotherapy in the treatment of advanced diffuse histiocytic lymphoma. Ann Intern Med. 1976;85(4):417–22. doi: 10.7326/0003-4819-85-4-417

12. Skarin AT, Rosenthal DS, Moloney WC, Frei E, 3rd. Combination chemotherapy of advanced non-Hodgkin lymphoma with bleomycin, adriamycin, cyclophosphamide, vincristine, and prednisone (BACOP). Blood. 1977;49(5):759–70. doi:

13. McKelvey EM, Gottlieb JA, Wilson HE, Haut A, Talley RW, Stephens R, et al. Hydroxyldaunomycin (Adriamycin) combination chemotherapy in malignant lymphoma. Cancer. 1976;38(4):1484–93. doi: 10.1002/1097-0142(197610)38:4<1484::aid-cncr2820380407>3.0.co;2-i

14. Fisher RI, Gaynor ER, Dahlberg S, Oken MM, Grogan TM, Mize EM, et al. Comparison of a standard regimen (CHOP) with three intensive chemotherapy regimens for advanced non-Hodgkin’s lymphoma. N Engl J Med. 1993;328(14):1002–6. doi: 10.1056/NEJM199304083281404

15. Gordon LI, Harrington D, Andersen J, Colgan J, Glick J, Neiman R, et al. Comparison of a second-generation combination chemotherapeutic regimen (m-BACOD) with a standard regimen (CHOP) for advanced diffuse non-Hodgkin’s lymphoma. N Engl J Med. 1992;327(19):1342–9. doi: 10.1056/NEJM199211053271903

16. Miller TP, Jones SE. Initial chemotherapy for clinically localized lymphomas of unfavorable histology. Blood. 1983;62(2):413–8. doi:

17. Miller TP, Jones SE, Rainey JM. Cyclophosphamide, doxorubicin, vincristine, and prednisone (CHOP) in patients with advanced Hodgkin’s disease: a Southwest Oncology Group Phase II Study. Cancer Treat Rep. 1983;67(7-8):739–40. doi:

18. Stashenko P, Nadler LM, Hardy R, Schlossman SF. Characterization of a human B lymphocyte-specific antigen. J Immunol. 1980;125(4):1678–85. doi:

19. Coiffier B, Haioun C, Ketterer N, Engert A, Tilly H, Ma D, et al. Rituximab (anti-CD20 monoclonal antibody) for the treatment of patients with relapsing or refractory aggressive lymphoma: a multicenter phase II study. Blood. 1998;92(6):1927–32. doi:

20. Maloney DG, Grillo-Lopez AJ, White CA, Bodkin D, Schilder RJ, Neidhart JA, et al. IDEC-C2B8 (Rituximab) anti-CD20 monoclonal antibody therapy in patients with relapsed low-grade non-Hodgkin’s lymphoma. Blood. 1997;90(6):2188–95. doi:

21. McLaughlin P, Grillo-Lopez AJ, Link BK, Levy R, Czuczman MS, Williams ME, et al. Rituximab chimeric anti-CD20 monoclonal antibody therapy for relapsed indolent lymphoma: half of patients respond to a four-dose treatment program. J Clin Oncol. 1998;16(8):2825–33. doi: 10.1200/JCO.1998.16.8.2825

22. Coiffier B, Lepage E, Briere J, Herbrecht R, Tilly H, Bouabdallah R, et al. CHOP chemotherapy plus rituximab compared with CHOP alone in elderly patients with diffuse large-B-cell lymphoma. N Engl J Med. 2002;346(4):235–42. doi: 10.1056/NEJMoa011795

23. Coiffier B, Thieblemont C, Van Den Neste E, Lepeu G, Plantier I, Castaigne S, et al. Long-term outcome of patients in the LNH-98.5 trial, the first randomized study comparing rituximab-CHOP to standard CHOP chemotherapy in DLBCL patients: a study by the Groupe d’Etudes des Lymphomes de l’Adulte. Blood. 2010;116(12):2040–5. doi: 10.1182/blood-2010-03-276246

24. Pfreundschuh M, Kuhnt E, Trumper L, Osterborg A, Trneny M, Shepherd L, et al. CHOP-like chemotherapy with or without rituximab in young patients with good-prognosis diffuse large-B-cell lymphoma: 6-year results of an open-label randomised study of the MabThera International Trial (MInT) Group. Lancet Oncol. 2011;12(11):1013–22. doi: 10.1016/S1470-2045(11)70235-2

25. Pfreundschuh M, Schubert J, Ziepert M, Schmits R, Mohren M, Lengfelder E, et al. Six versus eight cycles of bi-weekly CHOP-14 with or without rituximab in elderly patients with aggressive CD20+ B-cell lymphomas: a randomised controlled trial (RICOVER-60). Lancet Oncol. 2008;9(2):105–16. doi: 10.1016/S1470-2045(08)70002-0

26. Pfreundschuh M, Trumper L, Osterborg A, Pettengell R, Trneny M, Imrie K, et al. CHOP-like chemotherapy plus rituximab versus CHOP-like chemotherapy alone in young patients with good-prognosis diffuse large-B-cell lymphoma: a randomised controlled trial by the MabThera International Trial (MInT) Group. Lancet Oncol. 2006;7(5):379–91. doi: 10.1016/S1470-2045(06)70664-7

27. Tilly H, Morschhauser F, Sehn LH, Friedberg JW, Trneny M, Sharman JP, et al. Polatuzumab Vedotin in Previously Untreated Diffuse Large B-Cell Lymphoma. N Engl J Med. 2022;386(4):351–63. doi: 10.1056/NEJMoa2115304

28. Morschhauser F, Salles G, Sehn LH, Herrera AF, Friedberg JW, Trneny M, et al. Five-Year Outcomes of the POLARIX Study Comparing Pola-R-CHP and R-CHOP in Patients With Diffuse Large B-Cell Lymphoma. J Clin Oncol. 2025;43(35):3698–705. doi: 10.1200/JCO-25-00925

29. Neuendorff NR, Pugh K, Juthani R, Soudant C, Eochagain CM, Torka P. Inclusion, characteristics and reporting of older patients in recent registration trials for DLBCL: A systematic review. Blood Adv. 2025. doi: 10.1182/bloodadvances.2025017530

30. Abramson JS, McGree B, Noyes S, Plummer S, Wong C, Chen YB, et al. Anti-CD19 CAR T Cells in CNS Diffuse Large-B-Cell Lymphoma. N Engl J Med. 2017;377(8):783–4. doi: 10.1056/NEJMc1704610

31. Neelapu SS, Locke FL, Bartlett NL, Lekakis LJ, Miklos DB, Jacobson CA, et al. Axicabtagene Ciloleucel CAR T-Cell Therapy in Refractory Large B-Cell Lymphoma. N Engl J Med. 2017;377(26):2531–44. doi: 10.1056/NEJMoa1707447

32. Schuster SJ, Bishop MR, Tam CS, Waller EK, Borchmann P, McGuirk JP, et al. Tisagenlecleucel in Adult Relapsed or Refractory Diffuse Large B-Cell Lymphoma. N Engl J Med. 2019;380(1):45–56. doi: 10.1056/NEJMoa1804980

33. Wudhikarn K, Tomas AA, Flynn JR, Devlin SM, Brower J, Bachanova V, et al. Low toxicity and excellent outcomes in patients with DLBCL without residual lymphoma at the time of CD19 CAR T-cell therapy. Blood Adv. 2023;7(13):3192–8. doi: 10.1182/bloodadvances.2022008294

34. Cremasco F, Menietti E, Speziale D, Sam J, Sammicheli S, Richard M, et al. Cross-linking of T cell to B cell lymphoma by the T cell bispecific antibody CD20-TCB induces IFNgamma/CXCL10-dependent peripheral T cell recruitment in humanized murine model. PLoS One. 2021;16(1):e0241091. doi: 10.1371/journal.pone.0241091

35. Hutchings M, Morschhauser F, Iacoboni G, Carlo-Stella C, Offner FC, Sureda A, et al. Glofitamab, a Novel, Bivalent CD20-Targeting T-Cell-Engaging Bispecific Antibody, Induces Durable Complete Remissions in Relapsed or Refractory B-Cell Lymphoma: A Phase I Trial. J Clin Oncol. 2021;39(18):1959–70. doi: 10.1200/JCO.20.03175

36. Abramson JS, Ku M, Hertzberg M, Huang HQ, Fox CP, Zhang H, et al. Glofitamab plus gemcitabine and oxaliplatin (GemOx) versus rituximab-GemOx for relapsed or refractory diffuse large B-cell lymphoma (STARGLO): a global phase 3, randomised, open-label trial. Lancet. 2024;404(10466):1940–54. doi: 10.1016/S0140-6736(24)01774-4

37. Dickinson MJ, Carlo-Stella C, Morschhauser F, Bachy E, Corradini P, Iacoboni G, et al. Glofitamab for Relapsed or Refractory Diffuse Large B-Cell Lymphoma. N Engl J Med. 2022;387(24):2220–31. doi: 10.1056/NEJMoa2206913

38. Plonquet A, Haioun C, Jais JP, Debard AL, Salles G, Bene MC, et al. Peripheral blood natural killer cell count is associated with clinical outcome in patients with aaIPI 2-3 diffuse large B-cell lymphoma. Ann Oncol. 2007;18(7):1209–15. doi: 10.1093/annonc/mdm110

39. Gluck WL, Hurst D, Yuen A, Levine AM, Dayton MA, Gockerman JP, et al. Phase I studies of interleukin (IL)-2 and rituximab in B-cell non-hodgkin’s lymphoma: IL-2 mediated natural killer cell expansion correlations with clinical response. Clin Cancer Res. 2004;10(7):2253–64. doi: 10.1158/1078-0432.ccr-1087-3

40. Zöphel S, Küchler N, Jansky J, Hoxha C, Schäfer G, Weise JJ, et al. CD16+ as predictive marker for early relapse in aggressive B-NHL/DLBCL patients. Mol Cancer. 2024;23(1):210. doi: 10.1186/s12943-024-02123-7

41. Ehrhardt H, Schrembs D, Moritz C, Wachter F, Haldar S, Graubner U, et al. Optimized anti-tumor effects of anthracyclines plus Vinca alkaloids using a novel, mechanism-based application schedule. Blood. 2011;118(23):6123–31. doi: 10.1182/blood-2010-02-269811

42. Nakagawa J, Takahata T, Hyodo R, Chen Y, Hasui K, Sasaki K, et al. Evaluation for pharmacokinetic exposure of cytotoxic anticancer drugs in elderly patients receiving (R-)CHOP therapy. Sci Rep. 2021;11(1):785. doi: 10.1038/s41598-020-80706-2

43. Advani RH, Dickinson MJ, Fox CP, Kahl B, Herrera AF, Lenz G, et al. SKYGLO: A Global Phase III Randomized Study Evaluating Glofitamab Plus Polatuzumab Vedotin + Rituximab, Cyclophosphamide, Doxorubicin, and Prednisone (Pola-R-CHP) Versus Pola-R-CHP in Previously Untreated Patients with Large B-Cell Lymphoma (LBCL). Blood. 2024;144(Supplement 1):1718.1–.1. doi: 10.1182/blood-2024-194000

44. Falchi L, Offner F, de Vos S, Brody J, Morillo D, Linton K, et al. Fixed-duration epcoritamab + R-CHOP in patients with newly diagnosed DLBCL and high IPI scores (3–5) led to sustained remissions and disease-free survival beyond 3 years: Results from the EPCORE NHL-2 trial. Blood. 2025;146(Supplement 1):1955–. doi: 10.1182/blood-2025-1955

45. Michot J-M, Yagci M, Kargus K, Mohamed H, Chen J, Brouwer-Visser J, et al. Odronextamab plus chemotherapy in patients with previously untreated diffuse large B-cell lymphoma (DLBCL): First Results from part 1 of the Phase 3 Olympia-3 study. Blood. 2025;146(Supplement 1):65–. doi: 10.1182/blood-2025-65

46. Westin J, Fayad L, Steiner R, Ahmed S, Jain P, Malpica L, et al. primary analysis of the smart stop trial: Lenalidomide, tafasitamab, rituximab, and acalabrutinib alone and with combination chemotherapy in newly diagnosed diffuse large B-cell lymphoma. Blood. 2025;146(Supplement 1):477–. doi: 10.1182/blood-2025-477

47. Blaney SM, Boyett J, Friedman H, Gajjar A, Geyer R, Horowtiz M, et al. Phase I clinical trial of mafosfamide in infants and children aged 3 years or younger with newly diagnosed embryonal tumors: a pediatric brain tumor consortium study (PBTC-001). J Clin Oncol. 2005;23(3):525–31. doi: 10.1200/JCO.2005.06.544

48. Harahap Y, Samuel C, Anadalusia R, Syafhan NF. Analysis 4-Hydroxycyclophosphamide in Cancer Patients Plasma for Therapeutic Drug Monitoring of Cyclophosphamide. International Journal of Pharmacy and Pharmaceutical Sciences. 2016;8(9):7. doi: 10.22159/IJPPS.2016V8I9.12918

49. Joy MS, La M, Wang J, Bridges AS, Hu Y, Hogan SL, et al. Cyclophosphamide and 4-hydroxycyclophosphamide pharmacokinetics in patients with glomerulonephritis secondary to lupus and small vessel vasculitis. Br J Clin Pharmacol. 2012;74(3):445–55. doi: 10.1111/j.1365-2125.2012.04223.x

50. Kummerow C, Schwarz EC, Bufe B, Zufall F, Hoth M, Qu B. A simple, economic, time-resolved killing assay. Eur J Immunol. 2014;44(6):1870–2. doi: 10.1002/eji.201444518

51. Zheng S, Wang W, Aldahdooh J, Malyutina A, Shadbahr T, Tanoli Z, et al. SynergyFinder Plus: Toward Better Interpretation and Annotation of Drug Combination Screening Datasets. Genomics Proteomics Bioinformatics. 2022;20(3):587–96. doi: 10.1016/j.gpb.2022.01.004

52. Chapuy B, Wurm-Kuczera R, Michael R, Wang M, Pichler P, Huster A, et al. Phase II frontline chemolight R-pola-glo trial induces high and durable response rates in elderly and medically unfit/frail patients with aggressive B-cell lymphoma. Blood. 2025;146(Supplement 1):61–. doi: 10.1182/blood-2025-61

53. Peyrade F, Jardin F, Thieblemont C, Thyss A, Emile JF, Castaigne S, et al. Attenuated immunochemotherapy regimen (R-miniCHOP) in elderly patients older than 80 years with diffuse large B-cell lymphoma: a multicentre, single-arm, phase 2 trial. Lancet Oncol. 2011;12(5):460–8. doi: 10.1016/S1470-2045(11)70069-9

54. Tohda S, Sato T, Kogoshi H, Fu L, Sakano S, Nara N. Establishment of a novel B-cell lymphoma cell line with suppressed growth by gamma-secretase inhibitors. Leuk Res. 2006;30(11):1385–90. doi: 10.1016/j.leukres.2006.05.003

55. Amini RM, Berglund M, Rosenquist R, Von Heideman A, Lagercrantz S, Thunberg U, et al. A novel B-cell line (U-2932) established from a patient with diffuse large B-cell lymphoma following Hodgkin lymphoma. Leuk Lymphoma. 2002;43(11):2179–89. doi: 10.1080/1042819021000032917

56. Al-Katib AM, Smith MR, Kamanda WS, Pettit GR, Hamdan M, Mohamed AN, et al. Bryostatin 1 down-regulates mdr1 and potentiates vincristine cytotoxicity in diffuse large cell lymphoma xenografts. Clin Cancer Res. 1998;4(5):1305–14. doi:

57. Th’ng KH, Garewal G, Kearney L, Rassool F, Melo JV, White H, et al. Establishment and characterization of three new malignant lymphoid cell lines. Int J Cancer. 1987;39(1):89–93. doi: 10.1002/ijc.2910390116

58. Cox MC, Battella S, La Scaleia R, Pelliccia S, Di Napoli A, Porzia A, et al. Tumor-associated and immunochemotherapy-dependent long-term alterations of the peripheral blood NK cell compartment in DLBCL patients. Oncoimmunology. 2015;4(3):e990773. doi: 10.4161/2162402X.2014.990773

59. Luria SE, Delbruck M. Mutations of Bacteria from Virus Sensitivity to Virus Resistance. Genetics. 1943;28(6):491–511. doi: 10.1093/genetics/28.6.491

60. Law LW. Origin of the resistance of leukaemic cells to folic acid antagonists. Nature. 1952;169(4302):628–9. doi: 10.1038/169628a0

61. Gatenby RA, Silva AS, Gillies RJ, Frieden BR. Adaptive therapy. Cancer Res. 2009;69(11):4894–903. doi: 10.1158/0008-5472.CAN-08-3658

62. Coldman AJ, Goldie JH. A model for the resistance of tumor cells to cancer chemotherapeutic agents. Mathematical Biosciences. 1983;65(2):291–307. doi: 10.1016/0025-5564(83)90066-4

63. Law LW. Effects of Combinations of Antileukemic Agents on an Acute Lymphocytic Leukemia of Mice. Cancer Research. 1952;12(12):871–8. doi:

64. Ehrhardt H, Pannert L, Pfeiffer S, Wachter F, Amtmann E, Jeremias I. Enhanced anti-tumour effects of Vinca alkaloids given separately from cytostatic therapies. Br J Pharmacol. 2013;168(7):1558–69. doi: 10.1111/bph.12068

65. Kaschek L, Vialle J, Stopper G, Hoffmann MDA, Zophel S, Jansky J, et al. Fura-10, unlike fura-2, is suitable for long-term calcium imaging in natural killer (NK) cells without compromising cytotoxicity and can be combined with target cell death analysis. Cell Calcium. 2025;133:103091. doi: 10.1016/j.ceca.2025.103091

66. Knorck A, Marx S, Friedmann KS, Zophel S, Lieblang L, Hassig C, et al. Quantity, quality, and functionality of peripheral blood cells derived from residual blood of different apheresis kits. Transfusion. 2018;58(6):1516–26. doi: 10.1111/trf.14616

67. Pauck D, Picard D, Maue M, Taban K, Marquardt V, Blumel L, et al. An in vitro pharmacogenomic approach reveals subtype-specific therapeutic vulnerabilities in atypical teratoid/rhabdoid tumors (AT/RT). Pharmacol Res. 2025;213:107660. doi: 10.1016/j.phrs.2025.107660

68. Yadav B, Wennerberg K, Aittokallio T, Tang J. Searching for Drug Synergy in Complex Dose-Response Landscapes Using an Interaction Potency Model. Comput Struct Biotechnol J. 2015;13:504–13. doi: 10.1016/j.csbj.2015.09.001

