## Supplementary material for "R-CHOP in B cell non-Hodgkin lymphoma: balancing anti-tumor efficacy and NK-cell functionality": Jansky_Supplementary

Running title: R-CHOP effects on B-lymphoma cells, NK cell viability and cytotoxicity

**Key words:** non-Hodgkin lymphoma, diffuse large B-cell lymphoma (DLBCL), R-CHOP, vincristine, doxorubicin, cyclophosphamide, mafosfamide, rituximab, ADCC, natural killer cell, NK cell

A preprint of this manuscript has previously been published at BioRxiv (doi: 10.64898/2026.01.12.698321).

* Equal contribution

Markus Hoth and Eva C. Schwarz

Biophysics

Center for Integrative Physiology and Molecular Medicine (CIPMM)

Building 48

School of Medicine

Saarland University

66421 Homburg

Germany

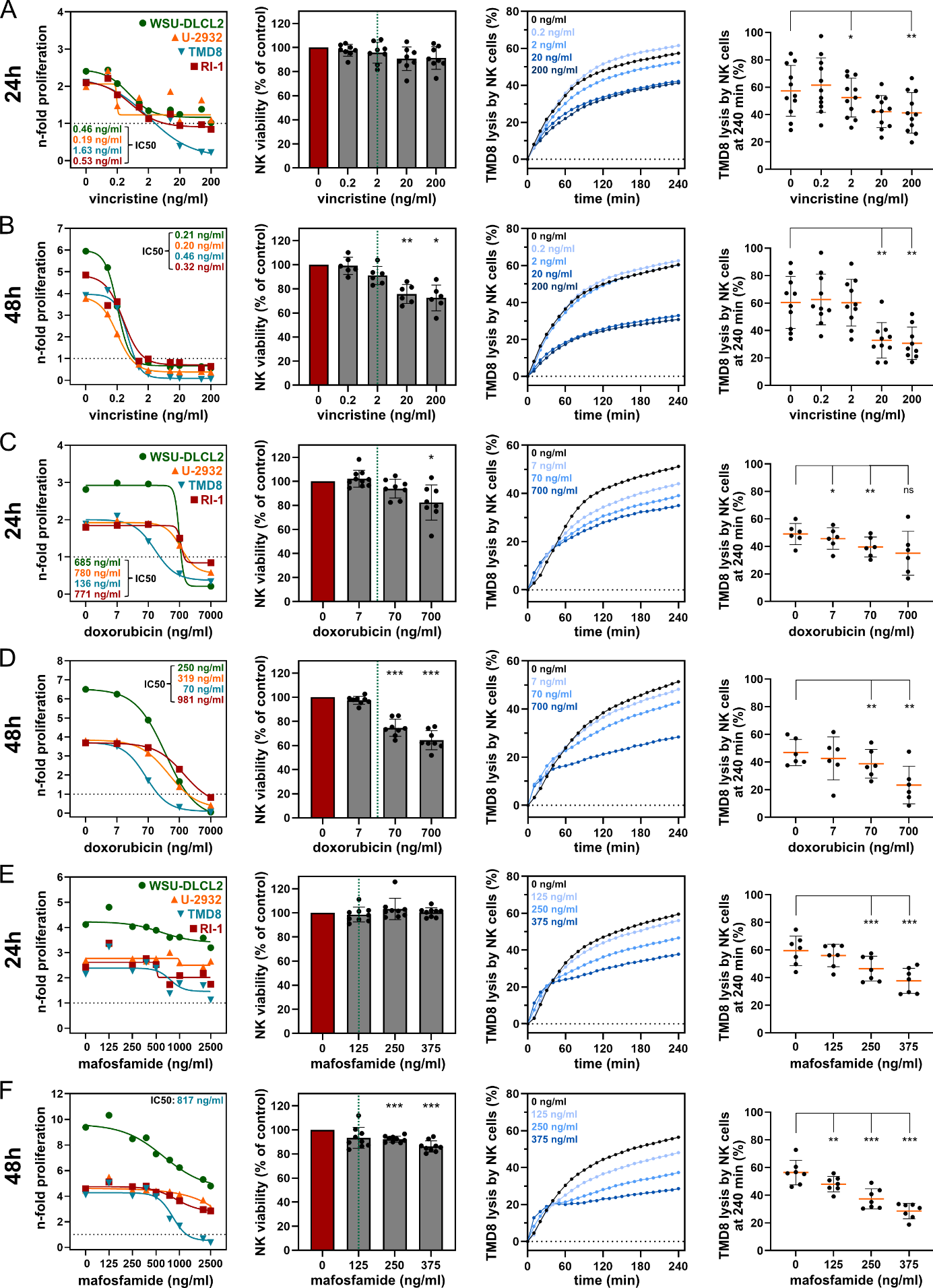

**Supplementary Figure 1: Vincristine, doxorubicin and mafosfamide dose-dependently reduce DLBCL cell proliferation, NK cell viability and cytotoxicity after 24- and 48-hour incubations.** (A and B) Vincristine, (C and D) doxorubicin and (E and F) mafosfamide incubations after 24 hours (A, C, E) and 48 hours (B, D, F). The first panels from left show n-fold proliferations of the DLBCL cell lines WSU-DLCL2 (green), U-2932 (orange), TMD8 (blue) and RI-1 (red). IC50 values (where calculable) are shown on the left panel side for 24h incubation and on the right panel side for 48h incubation and are color-coded according to cell line colors. Data points represent mean values (n=4-6). The second panels from left depict viabilities of expanded NK cells after incubation with different concentrations of vincristine, doxorubicin and mafosfamide, normalized to untreated control. Green dotted lines represent clinical concentrations. Bars represent mean ± SD (n=6-10). The third panels from left show mean TMD8 lysis kinetics with rituximab over 4 hours by expanded NK cells (E:T = 3:1) incubated with different concentrations of vincristine, doxorubicin and mafosfamide for 24h and 48h, as indicated by different shades of blue in the respective graph (n=6-11). The fourth panels from left depict the mean endpoint lysis ± SD of TMD8 with rituximab after 4 hours by expanded NK cells (E:T 3:1) incubated with different concentrations of vincristine, doxorubicin and mafosfamide for 24h and 48h (n=6-11). Asterisks highlight statistical differences. If no asterisks are shown, no statistical difference was detected.

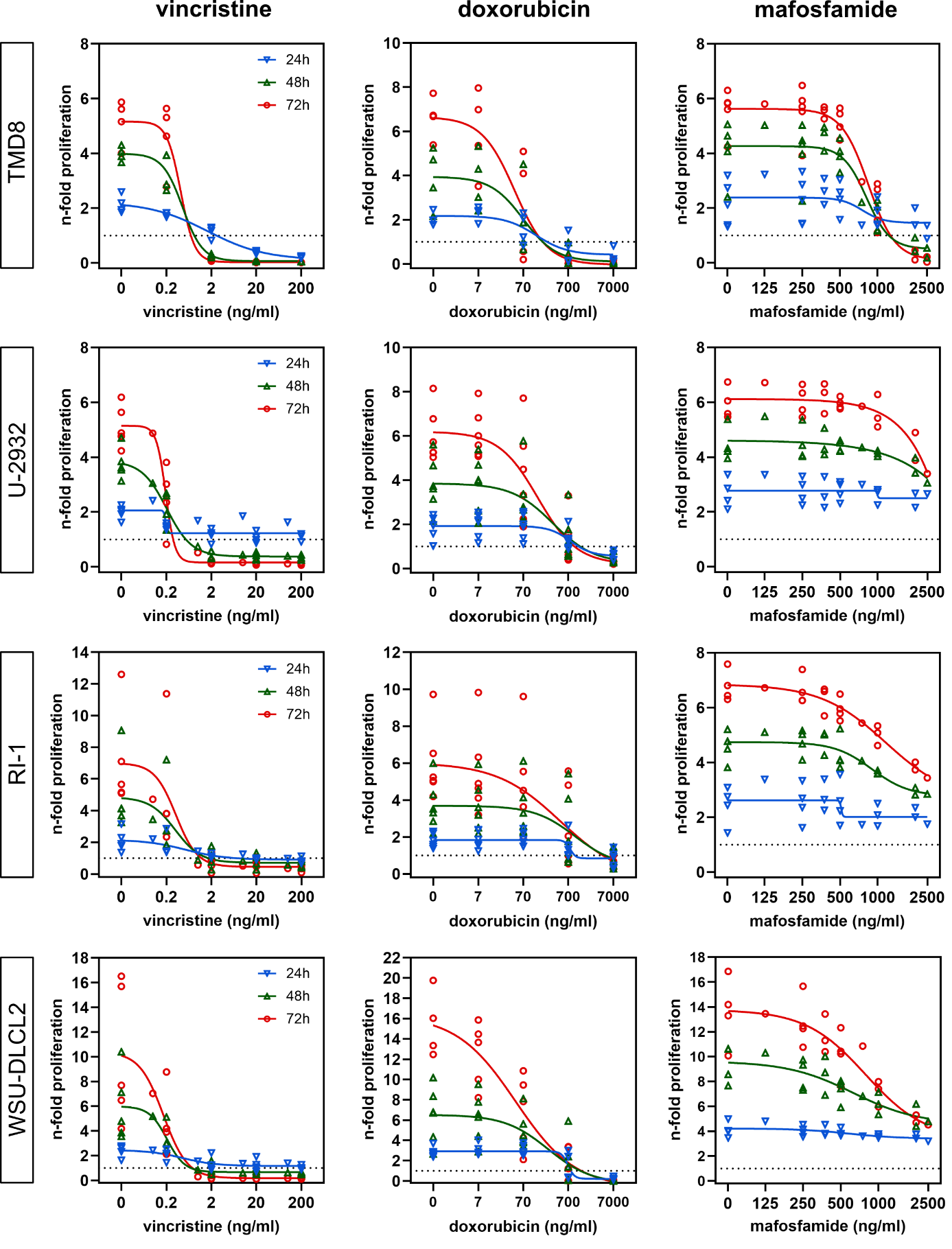

**Supplementary Figure 2:** **Dose-response curves of DLBCL cell lines treated with CHOP chemotherapeutics.** N-fold proliferation of the DLBCL cell lines TMD8, U-2932, RI-1 and WSU-DLCL2 (from top to bottom) after 24 (blue), 48 (green) and 72 hours (red) of incubation with different vincristine (left; n=4-5), doxorubicin (middle; n=4-6) or mafosfamide (right; n=4-5) concentrations. All data points of the experiments shown in Figure 2 A-C and Supplementary Figure 1 are included.

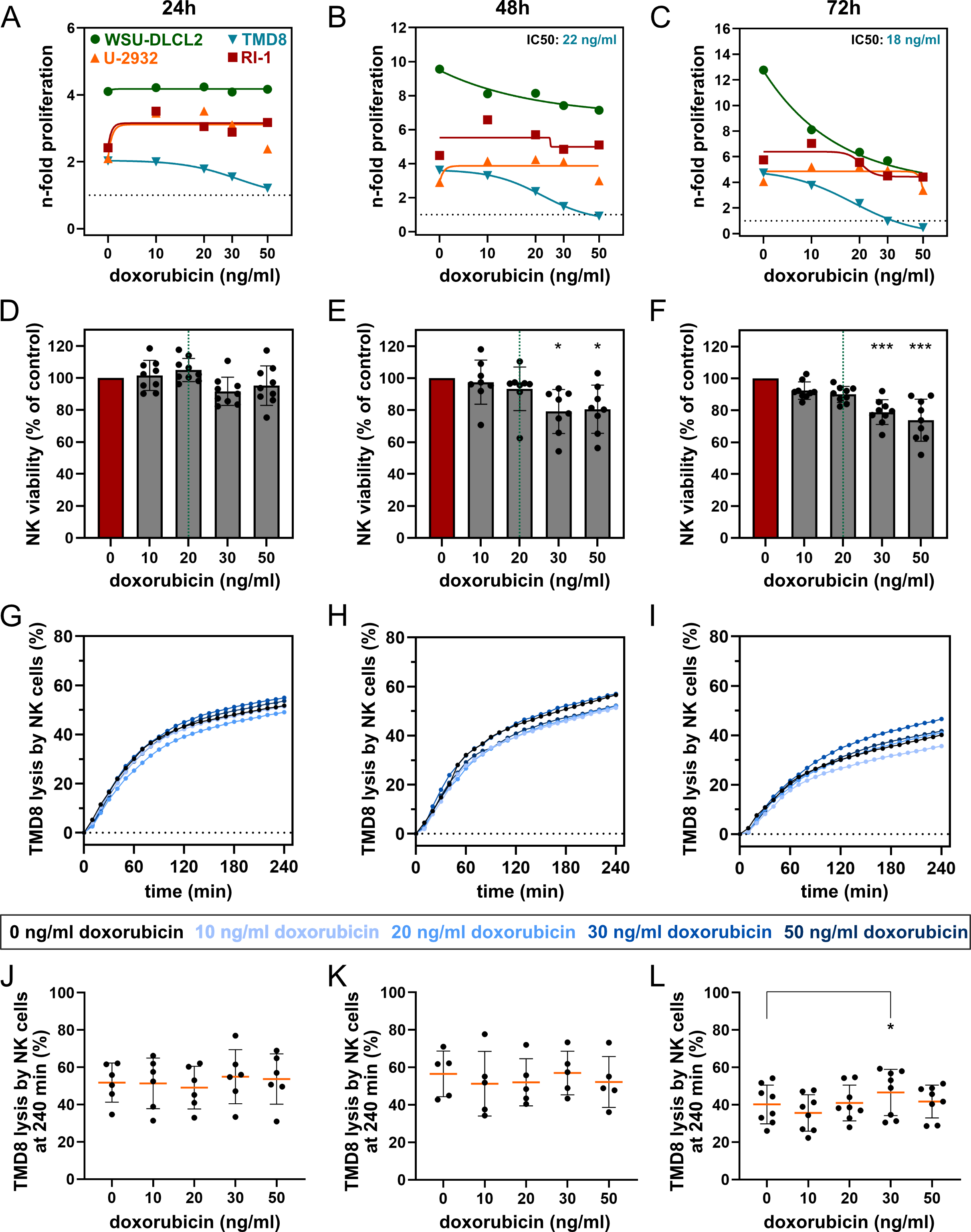

**Supplementary Figure 3: Low doxorubicin doses reduce NK cell viability, but not cytotoxicity, while barely affecting B cell proliferation.** (A-C) N-fold proliferation of the DLBCL cell lines WSU-DLCL2 (green), U-2932 (orange), TMD8 (blue) and RI-1 (red) after 24 (A), 48 (B) and 72 (C) hours of incubation with different doxorubicin concentrations. Where calculable, IC50 values are shown in the upper right (48h, 72h) corner and are color-coded according to cell line colors. Data points represent single values (n=1). (D-F) Viability of expanded NK cells after 24, 48 and 72 hours of incubation with different doxorubicin concentrations, normalized to untreated control. Green dotted lines represent clinical concentrations. Bars represent mean ± SD (n=8-9). (G-I) Mean TMD8 lysis kinetics over 4 hours by expanded NK cells (E:T 3:1) incubated with different doxorubicin concentrations (24h, 48h, 72h) as indicated by different shades of blue in the legend (n=5-8). (J-L) Mean endpoint lysis ± SD of TMD8 after 4 hours by expanded NK cells (E:T 3:1) incubated with different doxorubicin concentrations (24h, 48h, 72h, n=5-8).

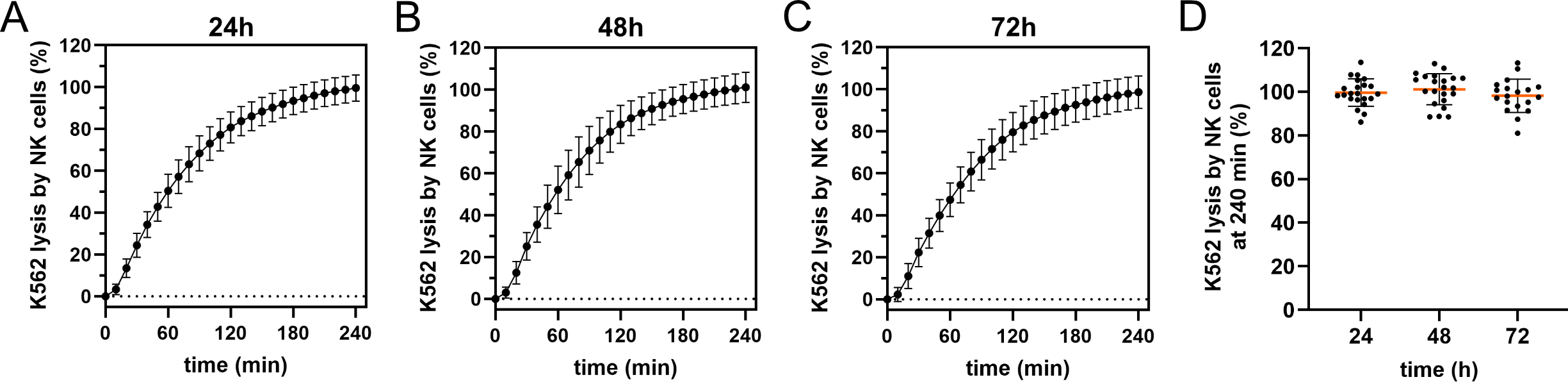

**Supplementary Figure 4: Untreated, expanded NK cells efficiently lyse K562 target cells after 24, 48 and 72 hours of incubation.** (A-C) Mean K562 lysis kinetics ± SD over 4 hours with untreated, control-incubated expanded NK cells (E:T 3:1; n=19-22). (A) K562 lysis kinetics after 24h of NK cell incubation. (B) K562 lysis kinetics after 48h of NK cell incubation. (C) K562 lysis kinetics after 72h of NK cell incubation. (D) Mean endpoint lysis ± SD of K562 after 4 hours with untreated, control-incubated (24h, 48h, 72h), expanded NK cells (E:T 3:1; n=19-22).

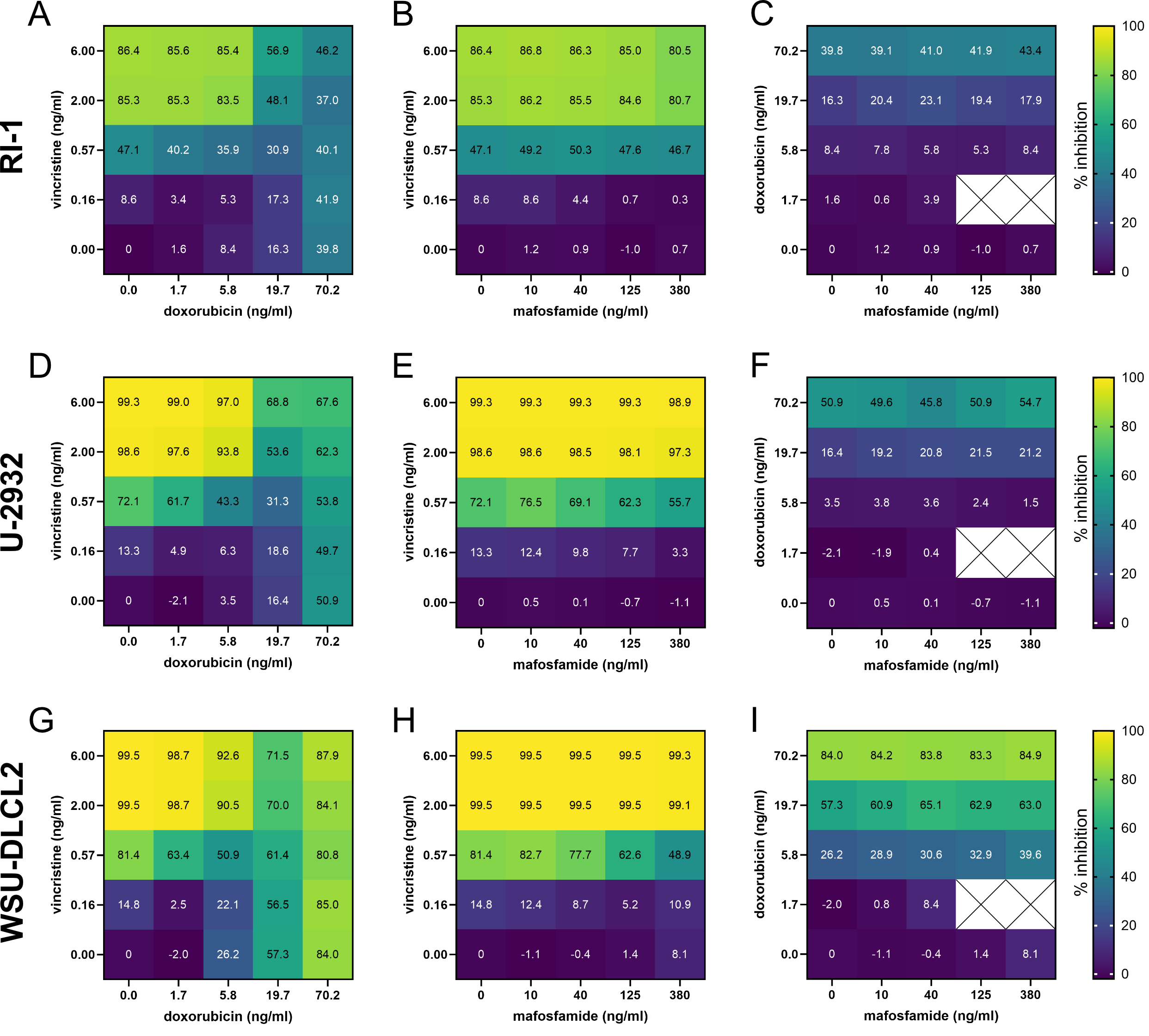

**Supplementary Figure 5:** **DLBCL line growths inhibition by two-drugs combinations.** Heat maps of the combinations of vincristine and doxorubicin (A, D, G), vincristine and mafosfamide (B, E, H) and doxorubicin and mafosfamide (C, F, I) for three different cell lines, RI-1 (upper row, A-C), U-2932 (middle row, D-F) and WSU-DLCL2 (lower row, G-I). Color code shows the percentage of inhibition. Cell viability was measured after 72 hours, and inhibition of growth was calculated.

| **RI-1** |  |
| --- | --- |
| 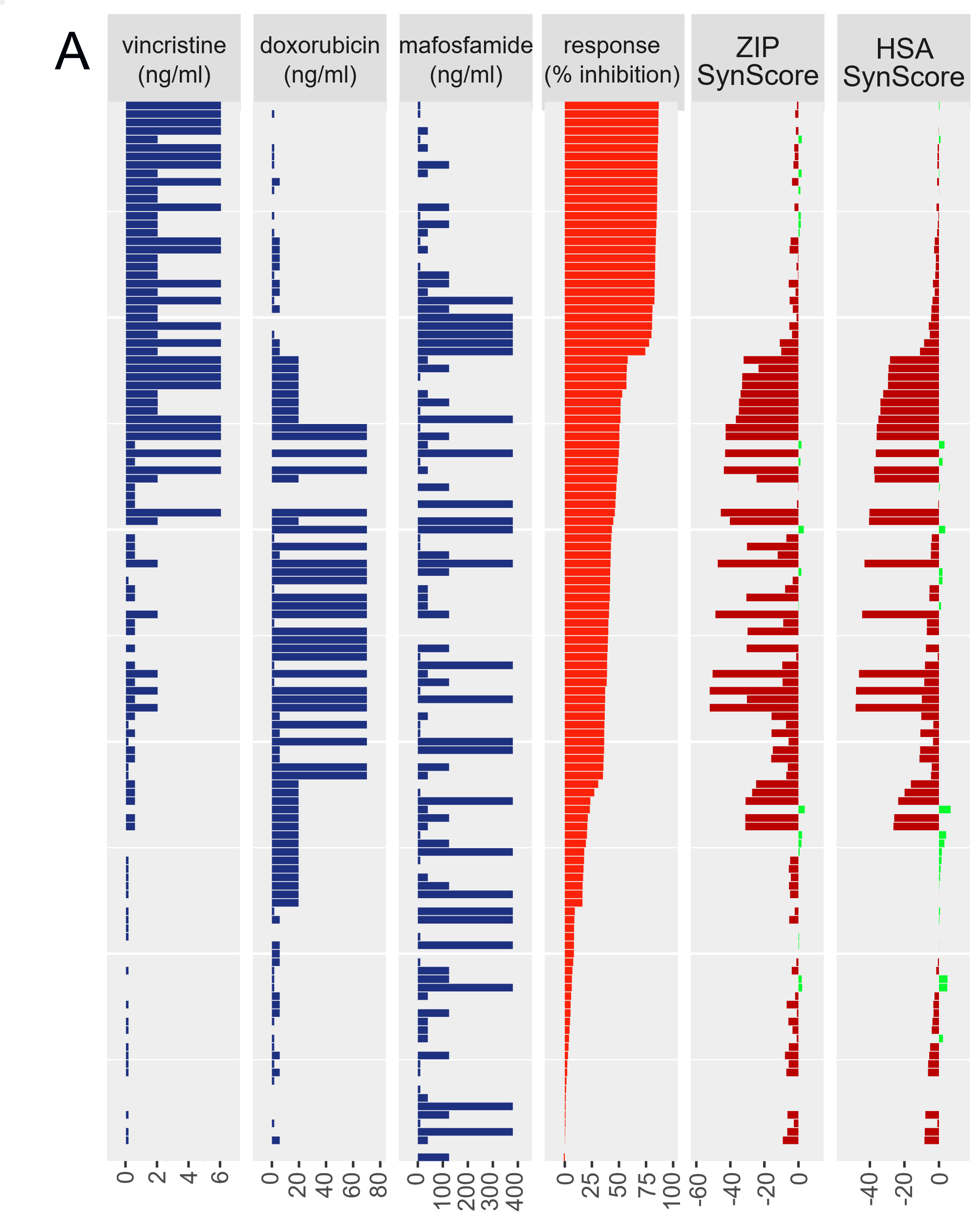 | 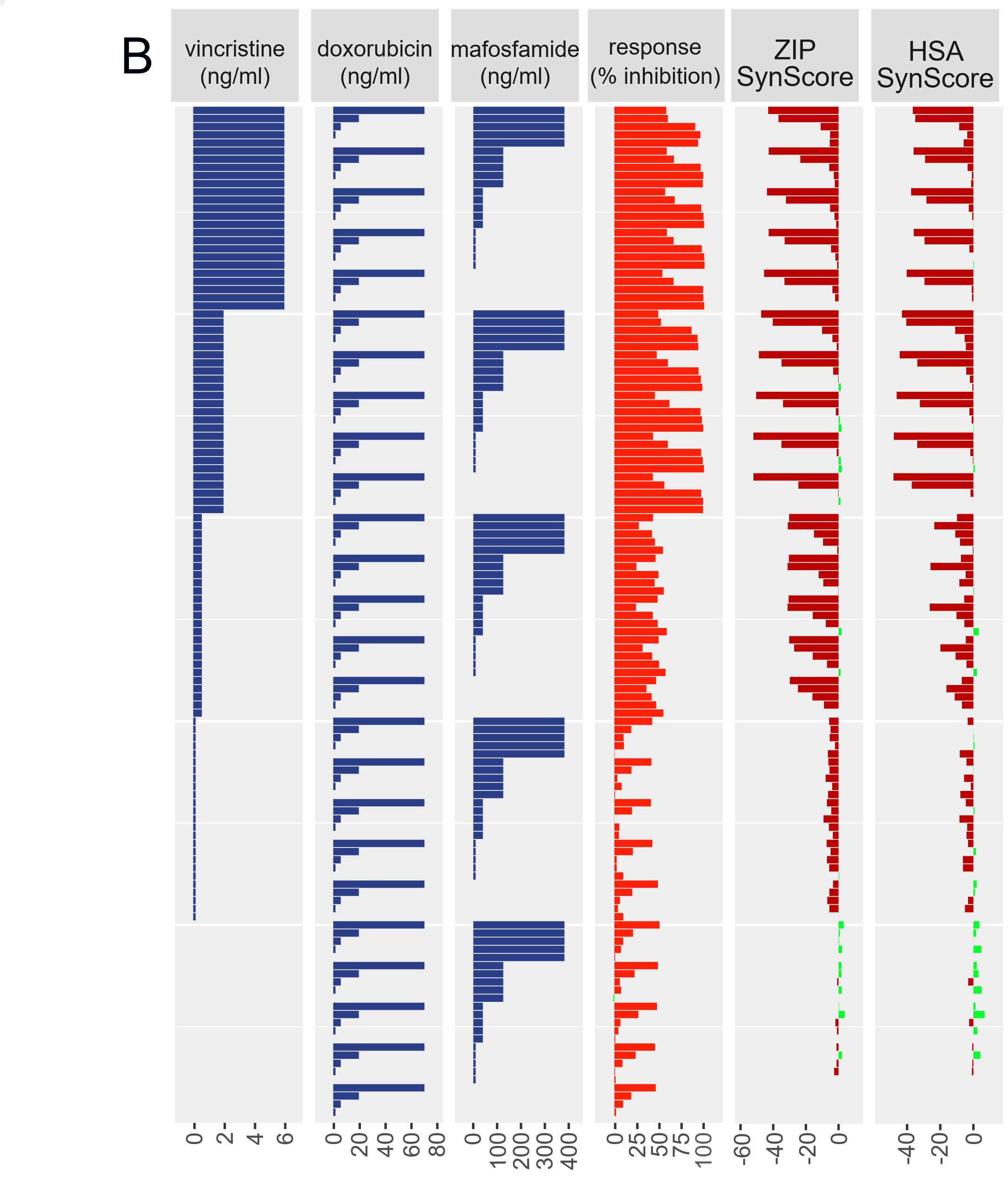 |
| **U-2932** |  |
| 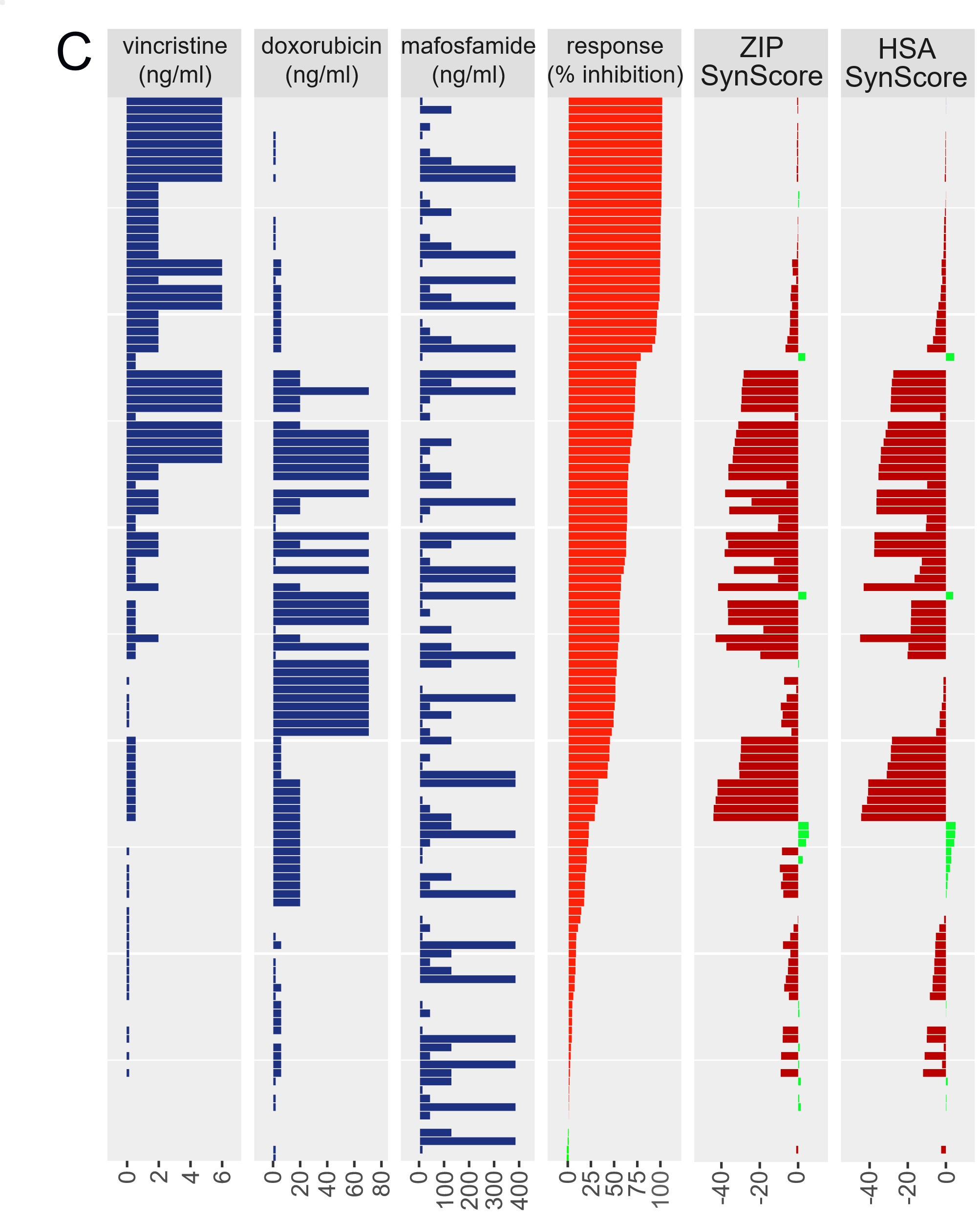 | 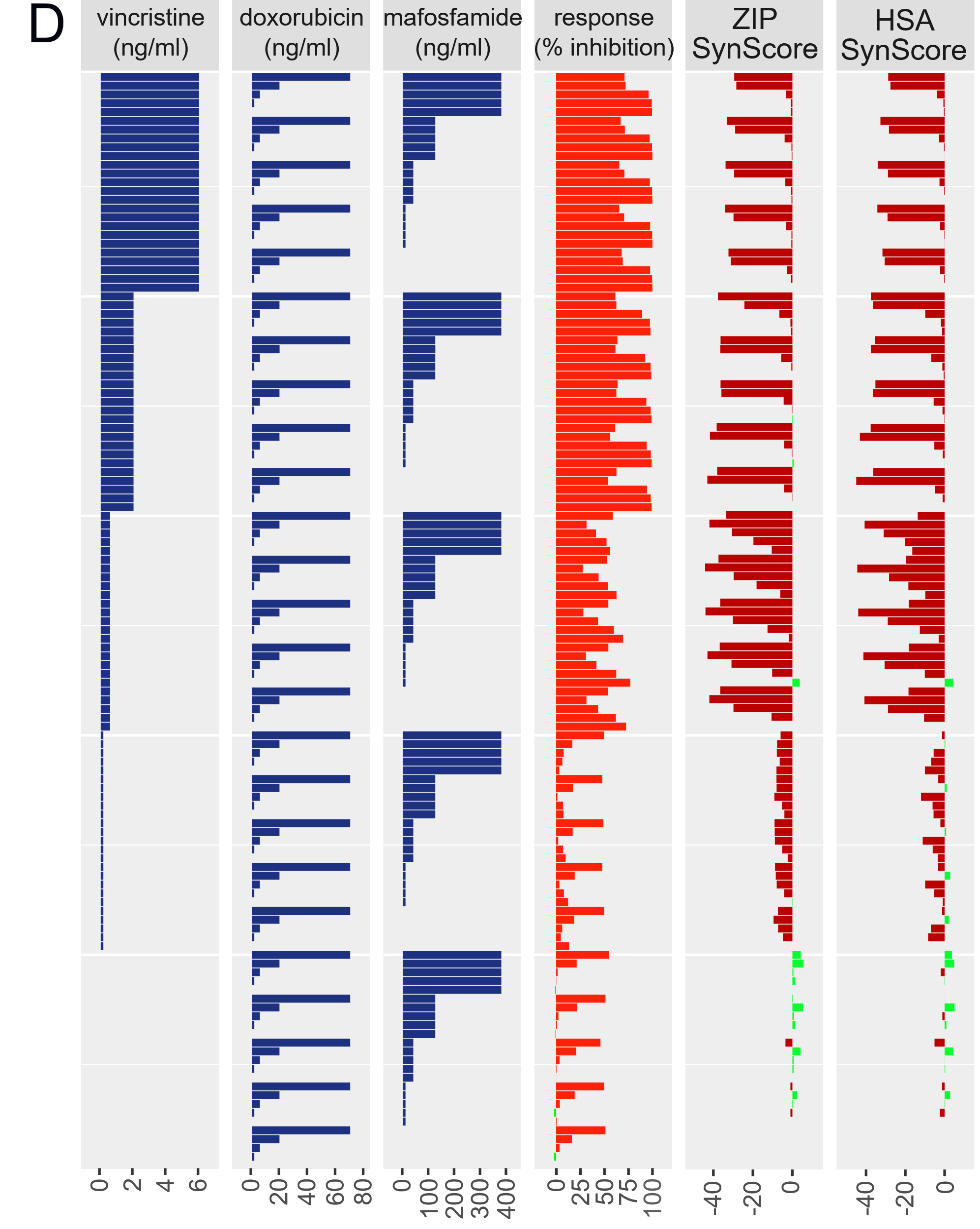 |
| **WSU-DCLC2** |  |

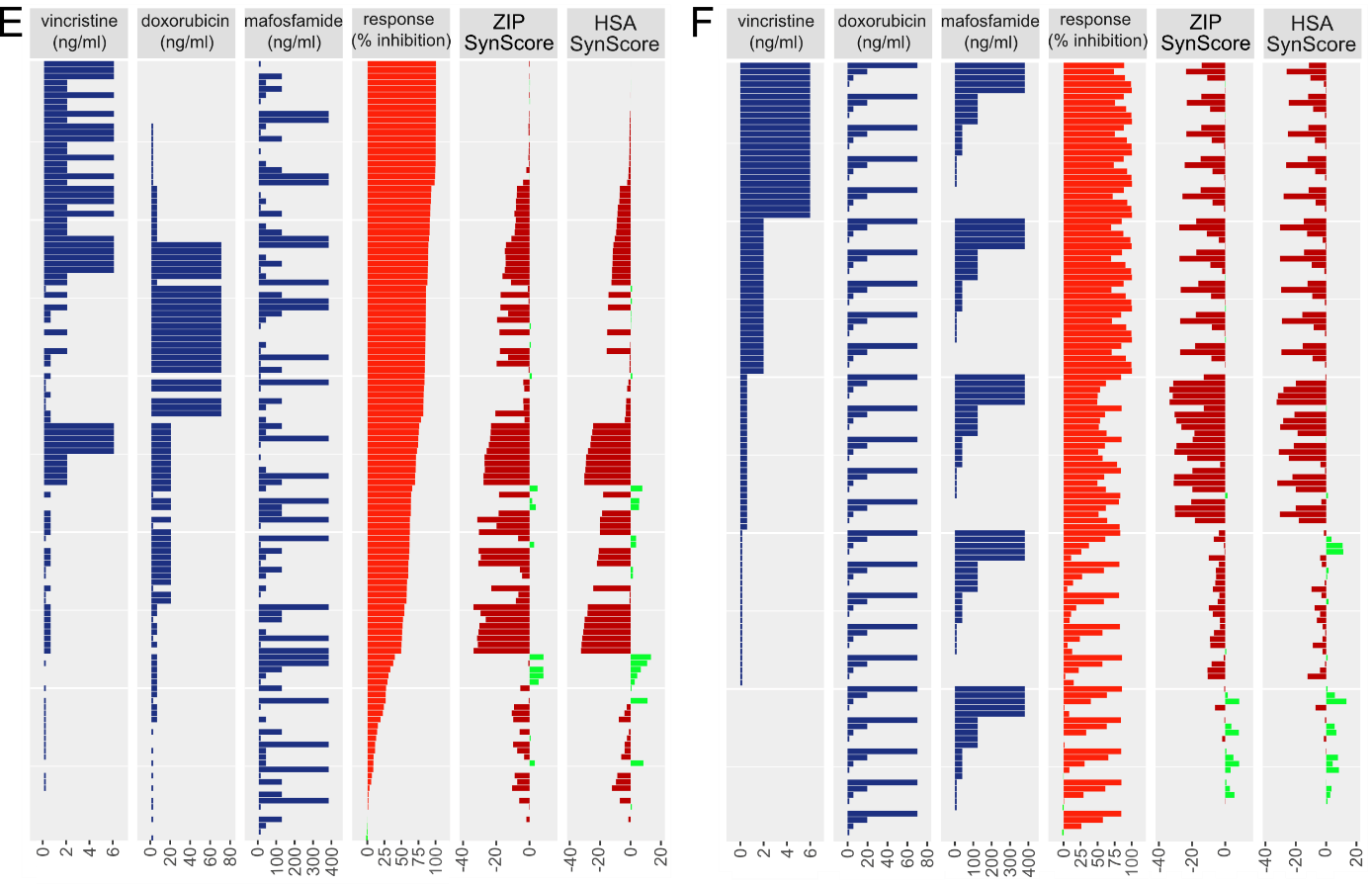

**Supplementary Figure 6: DLBCL cell line growth inhibition by two- and three-drug combination.** Synergy scores (ZIP and HSA) were calculated with Synergy Finder+ (https://synergyfinder.org/) for RI-1, U-2932 and WSU-DCLC2 cell lines for combinatorial concentrations of vincristine, doxorubicin and mafosfamide as given in Supplementary Figure 5 and Figure 3 A-C. Data are sorted for percentages of inhibition and for vincristine concentrations for RI-1 (A, B), U-2932 (C, D) and WSU-DCLC2 (E, F) cell lines.
